# Dysregulation of stress-induced translational control by *Porphyromonas gingivalis* in host cells

**DOI:** 10.1101/2022.01.24.477647

**Authors:** Alex A Knowles, Susan G Campbell, Neil A Cross, Prachi Stafford

## Abstract

Periodontitis, a chronic inflammatory gum disease, is caused in part by the periodontopathogen *Porphyromonas gingivalis*. Infection triggers activation of host inflammatory responses which induce stresses such as oxidative stress. Under such conditions, cells can activate the Integrated Stress Response (ISR), a signalling cascade which functions to determine cellular fate, by either downregulating protein synthesis and initiating a stress-response gene expression program, or if stress cannot be overcome, initiating programmed cell death. Recent studies have implicated the ISR signalling in both host antimicrobial defences and within the pathomechanism of certain microbes.

In this study, we investigated how *P. gingivalis* infection alters translation attenuation during oxidative stress-induced activation of the ISR pathway in oral epithelial cells. *P. gingivalis* infection alone did not result in ISR activation. In contrast, infection coupled with stress led to differential stress granule formation and composition, along with dysregulation of the microtubule network. Infection also heightened stress-induced translational repression, a response which could not be rescued by ISRIB, a potent ISR inhibitor. Heightened translational repression during stress was observed with both *P. gingivalis* conditioned media and outer membrane vesicles, implicating the role of a secretory factor, probably proteases known as gingipains, in this exacerbated translational repression. The effects of gingipain inhibitors and gingipains-deficient *P. gingivalis* mutants further confirmed these pathogen-specific proteases as the effector.

Gingipains are known to degrade the mammalian target of rapamycin (mTOR) and these studies implicate the gingipain-mTOR axis as the effector of host translational dysregulation during stress.

## INTRODUCTION

The oral cavity harbours a wide array of biofilm-forming bacteria, which form a symbiotic relationship with their host (1). However, in some cases the community becomes dysbiotic with an increased load of pathogenic bacteria, ultimately resulting in oral disease characterised by inflammation of gingival tissues (2, 3). In severe cases, disease progresses into the chronic condition known as periodontitis (3), the 6^th^ most prevalent disease worldwide affecting ∼743 million (4). Periodontal disease has been associated with a range of diseases including cardiovascular disease (5), rheumatoid arthritis (6), diabetes (7), cancer (8), Alzheimer’s disease (9) and Parkinson’s disease (10).

Periodontitis is caused by a variety of pathogenic bacteria, the most prominent pathogens being *Porphyromonas gingivalis*, the keystone pathogen, as well as *Tannerella forsythia* and *Treponema denticola* (2, 11). *P. gingivalis*’ invasion of oral epithelial cells disrupts intracellular homeostasis in several ways (12). One example is via the major virulence factor gingipains, extracellular cysteine proteases (13). These are known to degrade key host proteins, including the mammalian Target Of Rapamycin Complex 1 (mTORC1) (14, 15), which is central to many processes including protein synthesis and autophagy (16). In addition, *P. gingivalis* inhibits host antimicrobial and phagocytic responses, which can create a favourable replicative niche (12).

Progression of periodontitis leads to an increasingly cytotoxic environment within the periodontal pocket with increasing levels of bacterial metabolites and oxidative stress due to neutrophil activation (17). Under such stress conditions, host cells activate a number of signalling cascades, one of which is a concerted cellular reprogramming system, termed the Integrated Stress Response (ISR), which functions to determine cellular fate (18).

Functionally, the ISR initially causes a global down-regulation of protein synthesis, which sets out to conserve energy and allow the activation of a stress response gene expression program thereby allowing the cells to overcome the stress (18). A variety of stresses, including bacterial infection, activate one or more of four stress response kinases; Protein Kinase R (PKR), Protein Kinase R like ER Kinase (PERK), General Control Nondepressible 2 (GCN2) and Heme Regulated Inhibitor (HRI) (kinases reviewed by Donnelley *et al.* (19); bacteria and kinases reviewed in Knowles *et al.* (20)). Once activated, these stress response kinases converge upon the phosphorylation of the eukaryotic initiation factor 2 alpha subunit (eIF2α) at serine 51 (19, 21). eIF2α in its GTP bound form binds the initiator methionyl tRNA, forming the ternary complex, a prerequisite for functional translation initiation (22). During homeostatic translation, eIF2-GTP is hydrolysed to eIF2-GDP, following which eIF2-GTP is regenerated by eIF2B, allowing for subsequent rounds of translation initiation (23, 24). Stress-induced eIF2α phosphorylation blocks the ability of eIF2B to regenerate eIF2-GTP resulting in the abrogation of global translation by inhibiting the formation of active ternary complex (25–27). Translation may be stalled independently of eIF2α through the eIF4E binding protein 1 (4E-BP1) (28), regulated by mTORC1 (29, 30).

Independent of the upstream stimuli, translational shutoff pathways result in stalled messenger ribonucleoprotein particles (mRNPs), which are aggregated into cytoplasmic foci known as stress granules. These function to aid sorting of mRNPs into those which will be degraded, or re-initiation if stress is overcome and translation resumes (31, 32). Stress granules form within minutes and dissolve at a similar pace (33). Therefore, owing to the dynamic nature of their existence, ongoing retrograde transport of components along functioning microtubules is required (34).

In the context of infection, viruses have been well documented to dysregulate translational control and ISR function (35, 36). Recent studies have reported that bacterial species may also target the host translational control machinery and ISR function (Reviewed in Knowles *et al.* (20)). Several bacteria are known to activate host ISR stress response kinases upon infection including *Shigella flexneri*, *Salmonella* (37, 38), *Pseudomonas aeruginosa* (39), *Mycobacterium tuberculosis* (40), *Yersinia pseudotuberculosis* (41), Shiga toxin *Escherishia coli* (STEC) (42) and Group A *Streptococci* (43).

Downstream of ISR activation, *S. flexneri*, *Salmonella* and STEC induce stress granule formation during infection (37, 42). In the presence of exogenous stress, *E. coli* decreases the frequency of cells producing stress granules (44) whilst *S. flexneri* infection results in increased stress granule frequency with differing composition (45). The mechanism of stress granule modulation by *S. flexneri* is not fully elucidated but inhibition of mTORC1, whose function controls the motility of certain stress granule components, and dysregulation of the microtubule network have been proposed as a possible mechanism (38, 45).

Intracellular *P. gingivalis* have been shown to degrade mTOR in a manner dependent on lysine-specific gingipain, secreted by *P. gingivalis* (15). However, when secreted, both the lysine- and arginine-specific gingipains elicit the downregulation of mTOR activity acting through the PI3K-AKT pathway (46). Furthermore, *P. gingivalis* has been shown to induce activation of the Unfolded Protein Response (UPR) (47), which interlinks with the ISR (48). These findings together with the fact that periodontal infection produces possible stress through inflammation (12, 17) suggest that *P. gingivalis* infection may also manipulate the host translational control pathways and stress granule formation. The overall aim of this study was to determine whether *P. gingivalis* dysregulates host translational control during oxidative stress and alters stress granule dynamics.

## RESULTS

### *P. gingivalis* infection heightens translational repression and modulates stress granule formation during exogenous stress

Bacterial infection can lead to an oxidative stress environment which is known to activate the host integrated stress response. To determine if *P. gingivalis* can dysregulate the host ISR, the effect on protein synthesis in the presence and absence of sodium arsenite, a chemical inducer of oxidative stress, was monitored in infected H357 cells (t=2, 4 and 6h; MOI 1:00). While infection alone did not induce the ISR (Fig S1A-C), a heightened stress-induced translational inhibition of 0.5-fold was observed when cells were treated with both *P. gingivalis* and oxidative stress (Fig 1A). As this increased inhibition of translation was observed at all infection time points, further studies were conducted after 2h infection.

**FIG 1.**
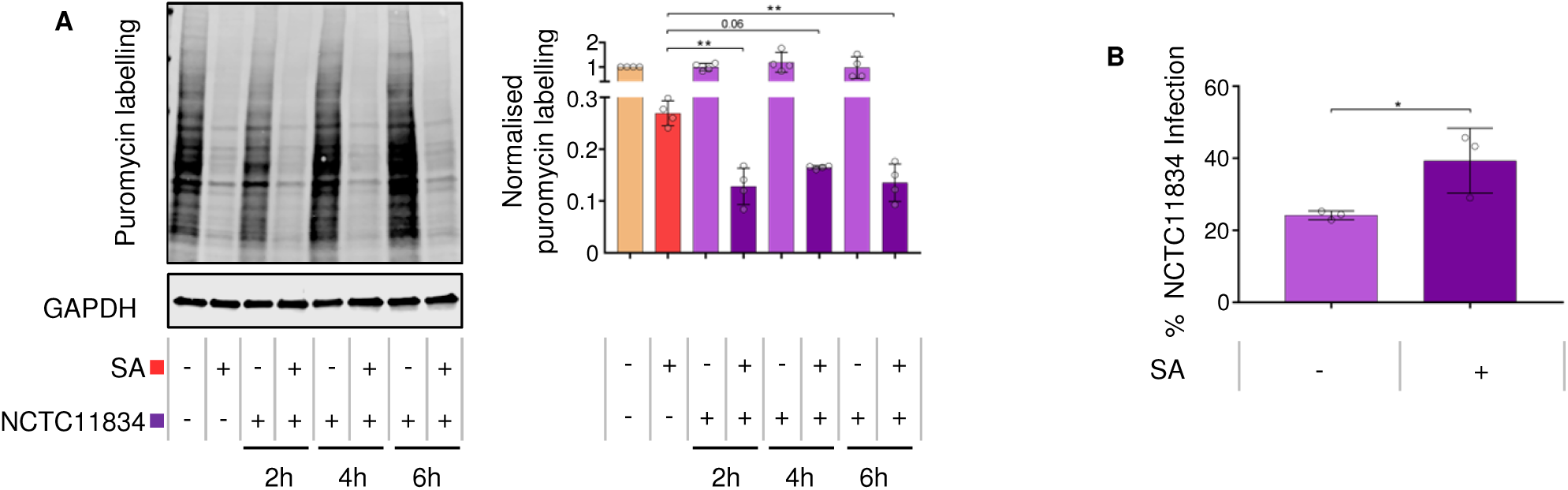
*P. gingivalis* infection heightens translational repression during oxidative stress. (A) Relative rate of protein synthesis in infected H357 cells following puromycin uptake (left) and relative quantification by first normalised to GAPDH and then to untreated sample (mean ± SD, n=4). (B) Percentage of H357 cells displaying internalised antibody signal for *P. gingivalis* post-infection after two hours (mean ± SD, n=3). **, *P* ≤ 0.01; *, *P* ≤ 0.05 according to Kruskal-Wallis with Conover-Inman *post-hoc*.

To determine whether oxidative stress influences *P. gingivalis* invasion, the percentage of cells infected with *P. gingivalis* in the presence or absence of sodium arsenite was quantified. In the absence of oxidative stress *P. gingivalis* infected 24% of cells compared to 39% of total cells in the presence of oxidative stress (Fig 1B).

To establish whether *P. gingivalis* infection could impact the formation of stress granules, the number of stress granules was quantified in cells displaying internalised *P. gingivalis* (Fig S1D). Cells treated with oxidative stress induced on average the formation of 36.2 stress granules per cell with an average size area of 2.25µM^2^. Within the bacteria treated population, neither uninfected nor infected cells showed evidence of stress granules (Fig 2A). In contrast, when *P. gingivalis* infection was coupled with oxidative stress, the frequency of stress granules increased on average to 59.4 per cell (Fig 2B). Differences between uninfected and infected cells within this population were further characterised and a decrease in stress granules frequency was observed in uninfected cells with the average area (2.2µM**^2^**) showing slight variance (Fig 2B).

**FIG 2.**
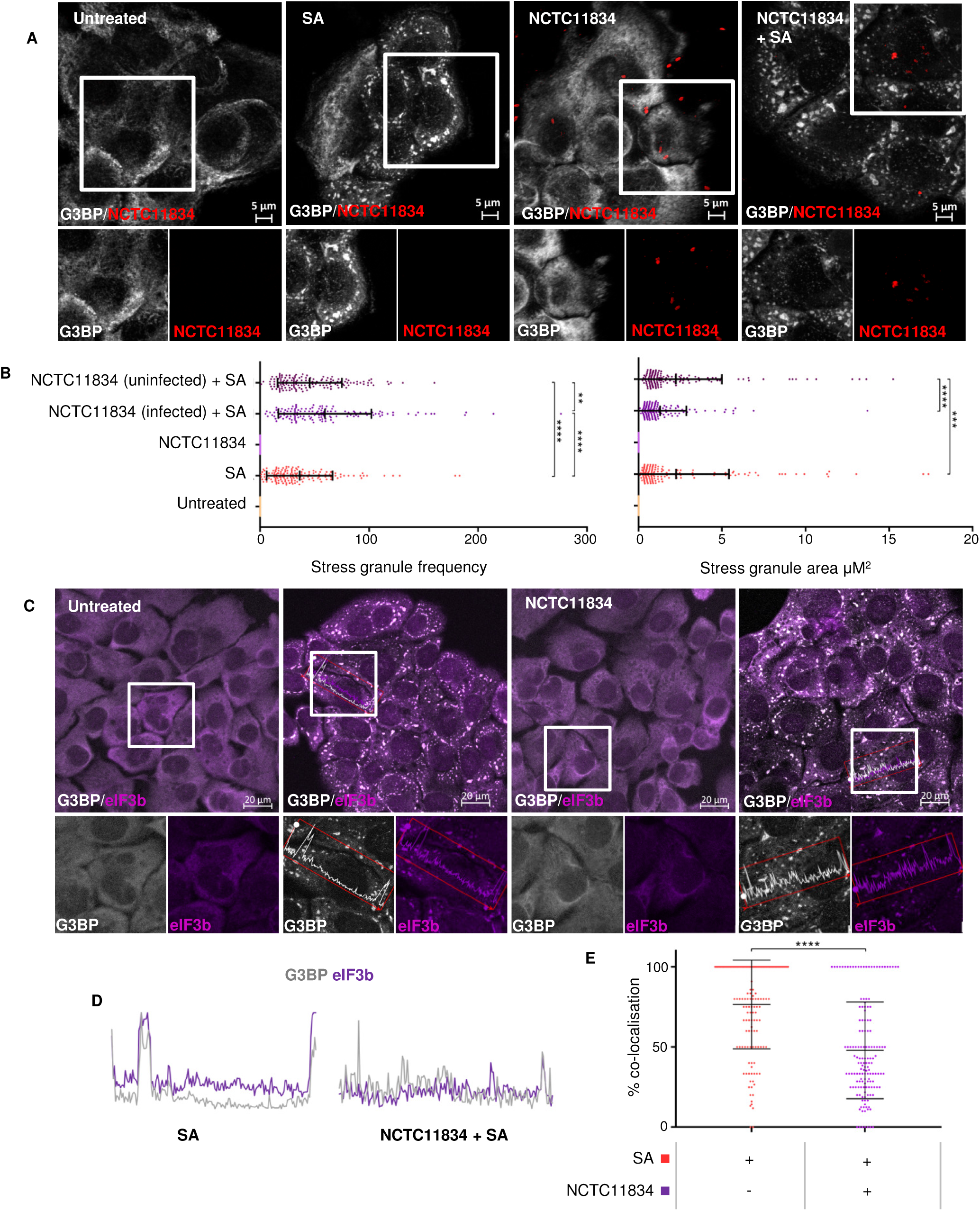
*P. gingivalis* infection modulates stress granule formation during oxidative stress in H357 cells. (A) Stress granule formation as visualised by G3BP1 (white) and *P. gingivalis* (red) using confocal microscopy and Z-stacks following bacterial challenge in the presence or absence of sodium arsenite. (B) Average area and frequency of stress granules determined in host cells (n=3, 50 cells per biological replicate). (C) Co-localisation of G3BP1 (white) and eIF3B (purple) in stress granules, as assessed by immunofluorescence. (D) Representative line segments of colour profiles taken from H357 cells challenged with sodium arsenite with or without *P. gingivalis* infection, where intensity peaks correspond to stress granules. (E) Percentage of colocalisation of eIF3b and G3BP1 (n=3, 50 cells per biological replicate). **** *P* ≤ 0.001; ***, *P* ≤ 0.001; **, *P* ≤ 0.01 according to Kruskal-Wallis with Conover-Inman *post-hoc*.

As stress granule composition is known to be stress dependent (49), the localisation of eIF3b and G3BP in stress granules was analysed (Fig 2C). During oxidative stress, eIF3b colocalised highly with G3BP positive stress granules (Fig 2D) (mean 75%). However, in the presence of *P. gingivalis* and oxidative stress, the mean percentage colocalization of eIF3b to G3BP decreased to 50% (Fig 2D). Given that *P. gingivalis* is known to degrade several host proteins, the potential for both G3BP or eIF3b degradation was investigated using immunoblotting. No degradation was observed (Fig S2), thereby suggesting the ability of *P. gingivalis* to modulate host stress granule frequency and composition.

### *P. gingivalis* heightens translational repression independently of eIF2α

Translational stalling during stress is classically mediated via the phosphorylation of alpha subunit of eIF2 at serine 51 (19, 21). The relative levels of total and p-eIF2a in *P. gingivalis* infected cells treated with or without oxidative stress were determined by immunoblotting (Fig 3A). Similar basal level of p-eIF2a (Fig 3A) was observed in *P. gingivalis* infected cells and the untreated control. Strikingly, despite the increased translational repression observed when *P. gingivalis* infection was co-treated with oxidative stress, a decrease in levels of p-eIF2a was observed compared to the oxidative stress-only treatment (Fig 3A).

**FIG 3.**
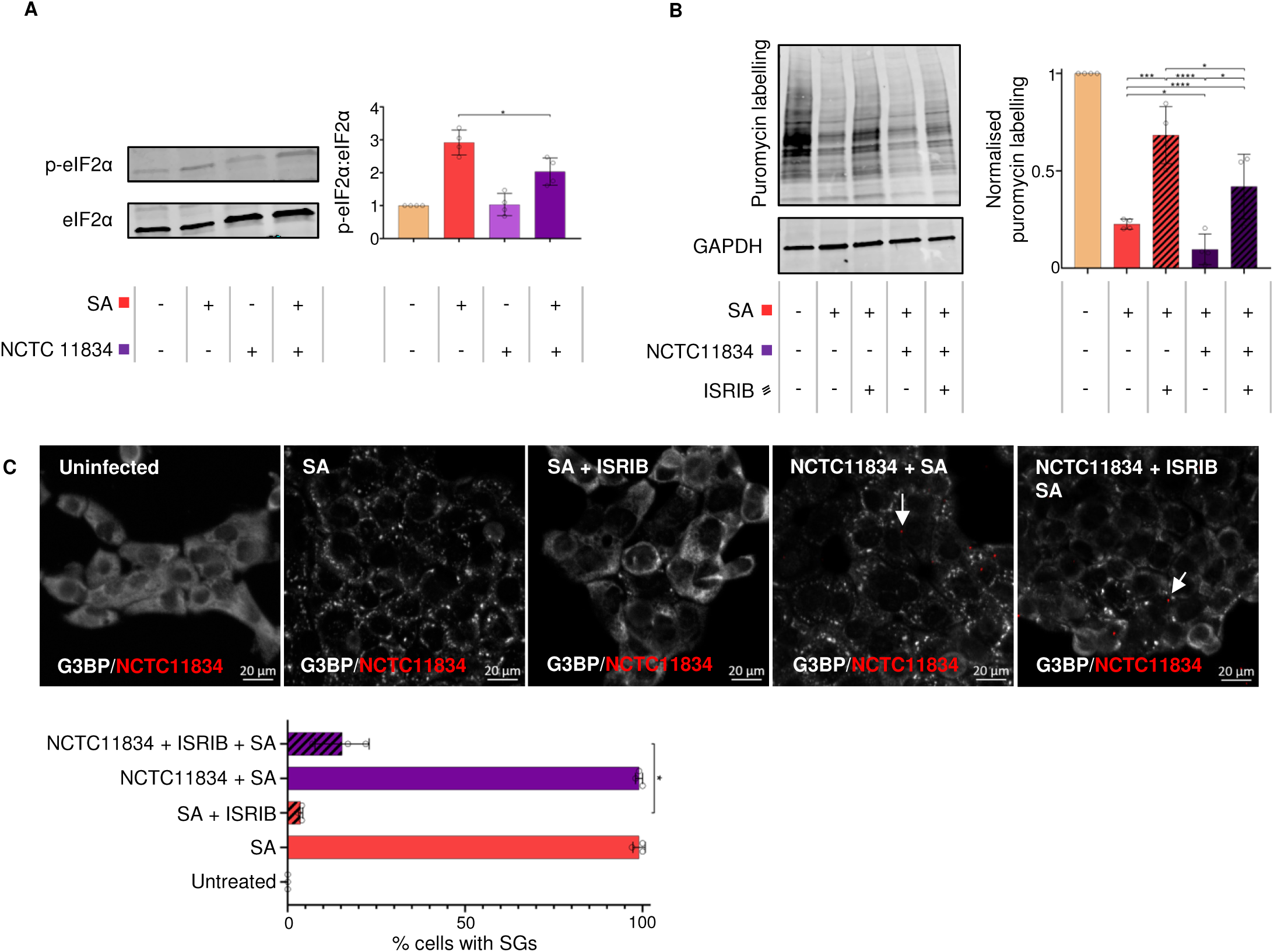
*P. gingivalis* heightens translational repression independently of eIF2α in the presence of stress. (A) Level of total and phosphorylated eIF2a (left) and ratio of phosphorylated eIF2a to total eIF2a after bacterial treatment of H357 cells in the presence or absence of sodium arsenite as determined by immunoblotting (right) (mean ± SD, n=4). (B) Relative rate of protein synthesis when H357 cells were treated as above and with ISRIB as determined by puromycin uptake (left) and the concentration relative to GAPDH (right) (mean ± SD, n=4). (C) Stress granule (SG) formation as visualisation of G3BP1 (white) and *P. gingivalis* (red) using confocal microscopy and Z-stacks (n=3, 100 cells per biological replicate). **** *P* ≤ 0.001; ***, *P* ≤ 0.001; **, *P* ≤ 0.01; *, *P* ≤ 0.05 according to Kruskal-Wallis with Conover-Inman *post-hoc*.

The small molecular ISR Inhibitor (ISRIB) has been shown to reverse the effects of p-eIF2a on translational inhibition and stress granule formation (50). Therefore, experiments were carried out to determine whether ISRIB could attenuate the heightened translational repression and modulation of SG formation during *P. gingivalis* infection during oxidative stress. In keeping with previous studies during ISRIB treatment alone, protein synthesis remained at steady state rates (Fig S3). In the presence of *P. gingivalis* and oxidative stress, ISRIB was only able to rescue translation 0.09-fold compared to 0.5-fold rescue during oxidative stress (Fig 3B). When the frequency of stress granules containing cells was quantified, ISRIB inhibited oxidative stress-induced stress granule formation 0.95-fold compared to only 0.84-fold in the presence of *P. gingivalis* and oxidative stress (Fig 3C). Collectively these data suggest that the heightened translational repression is independent of the ISR and cannot be rescued by ISRIB.

### *P. gingivalis* heightens translational repression via the action of a secretory factor

As uninfected cells within the infected population displayed increased stress granule frequency during oxidative stress and infection, the effect of *P. gingivalis* conditioned media was investigated to determine whether the observed effects were due to secreted bacterial components. Cells treated with conditioned media and oxidative stress decreased translation 0.2-fold compared to the oxidative stress only treatment (Fig 4A). Similar to the bacterial infection, cells treated with *P. gingivalis* conditioned media and oxidative stress decreased the levels of p-eIF2a compared to oxidative stress only treatment (Fig 4B). Taken together, these findings demonstrate that factors released by *P. gingivalis* are capable of heightening oxidative stress-induced translational inhibition.

**FIG 4.**
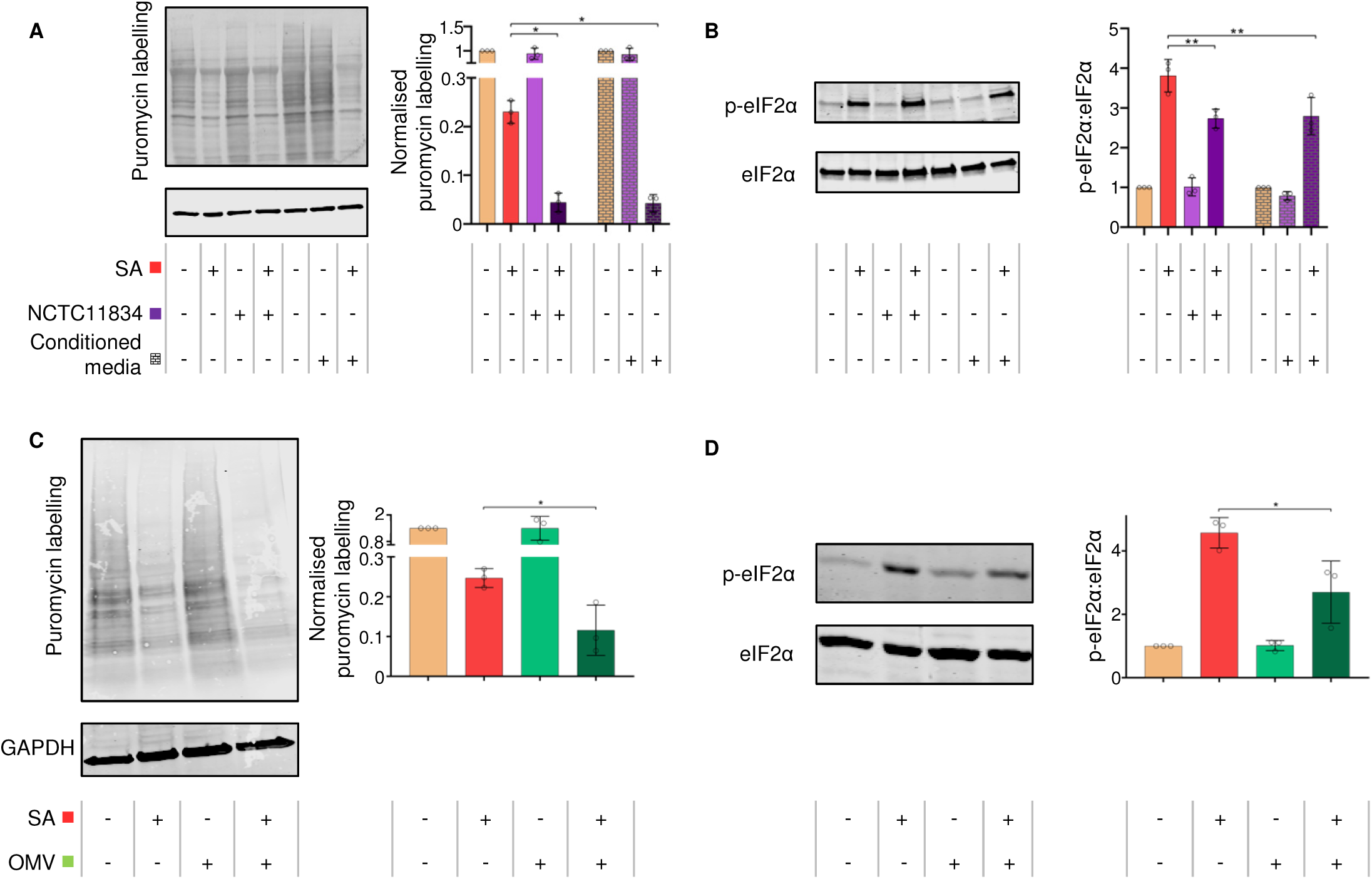
*P. gingivalis* heightens translational repression via the action of a secretory factor. (A) Relative rate of protein synthesis as determined by puromycin uptake (left) and quantification relative to GAPDH (right), when H357 cells were treated with filtered conditioned media recovered from cells previously treated with *P. gingivalis* (NCTC11834) in the presence or absence of sodium arsenite. (B) Levels of phosphorylated eIF2α (left) and concentration of phosphorylated to total eIF2α (right) when probed using immunoblotting (mean ± SD, n=3). (C) H357 cells were challenged with purified *P. gingivalis* OMV vesicles (100 µg/ml, t=2h) with or without sodium arsenite for the final 30 min and the relative rate of protein synthesis measured by puromycin uptake (left) and concentration relative to GAPDH (right) and (D) the levels of phosphorylated eIF2α (left) and the concentration of phosphorylated to total eIF2α (right) with OMVs were probed using immunoblotting (mean ± SD, n=3). **, *P* ≤ 0.01; *, *P* ≤ 0.05 according to Kruskal-Wallis with Conover-Inman *post-hoc*.

To establish which secreted bacterial constituents elicited the heightened translational inhibition observed during stress, cells were challenged with *P. gingivalis* outer membrane vesicles (OMVs) or purified lipopolysaccharide (LPS). OMVs (1µg/mL, 10µg/mL and 100µg/mL; t=2h) did not induce stress (Fig S4A&B). In the presence of oxidative stress, OMVs (100µg/mL, t=2h) heightened translational repression 0.53-fold (Fig 4C) and decreased p-eIF2a 0.41-fold (Fig 4D).

Purified LPS (1, 5 and 10µg/mL, t=2h) derived from *P. gingivalis* NCTC11834 did not induce stress (Fig S4C&D). In the presence of oxidative stress, LPS (10µg/mL, t=2h) did not alter translational repression or p-eIF2a (Fig S4E&F). This indicates that the heightened translational repression induced by *P. gingivalis* can be attributed to a secretory component distinct from LPS but present within the OMV fractions isolated.

### *P. gingivalis* dysregulates mTOR signalling during stress

Upon stress, mTORC1 has also been shown to contribute to translational control (51). Previously, *P. gingivalis* has been shown to both inhibit and degrade mTORC1 through the activity of its gingipains (15, 46). Given that the heightened translational repression during *P. gingivalis* infection and oxidative stress was independent of eIF2a signalling, the role of mTORC1 was evaluated. The effect of the selective mTOR inhibitor, rapamycin (400nM, t=1h) on mTORC1 during oxidative stress was determined. Similar to *P. gingivalis* infection, rapamycin, in the presence of oxidative stress, heightened translational repression 0.36-fold (Fig 5A), which was independent of p-eIF2a (Fig 5B).

**FIG 5.**
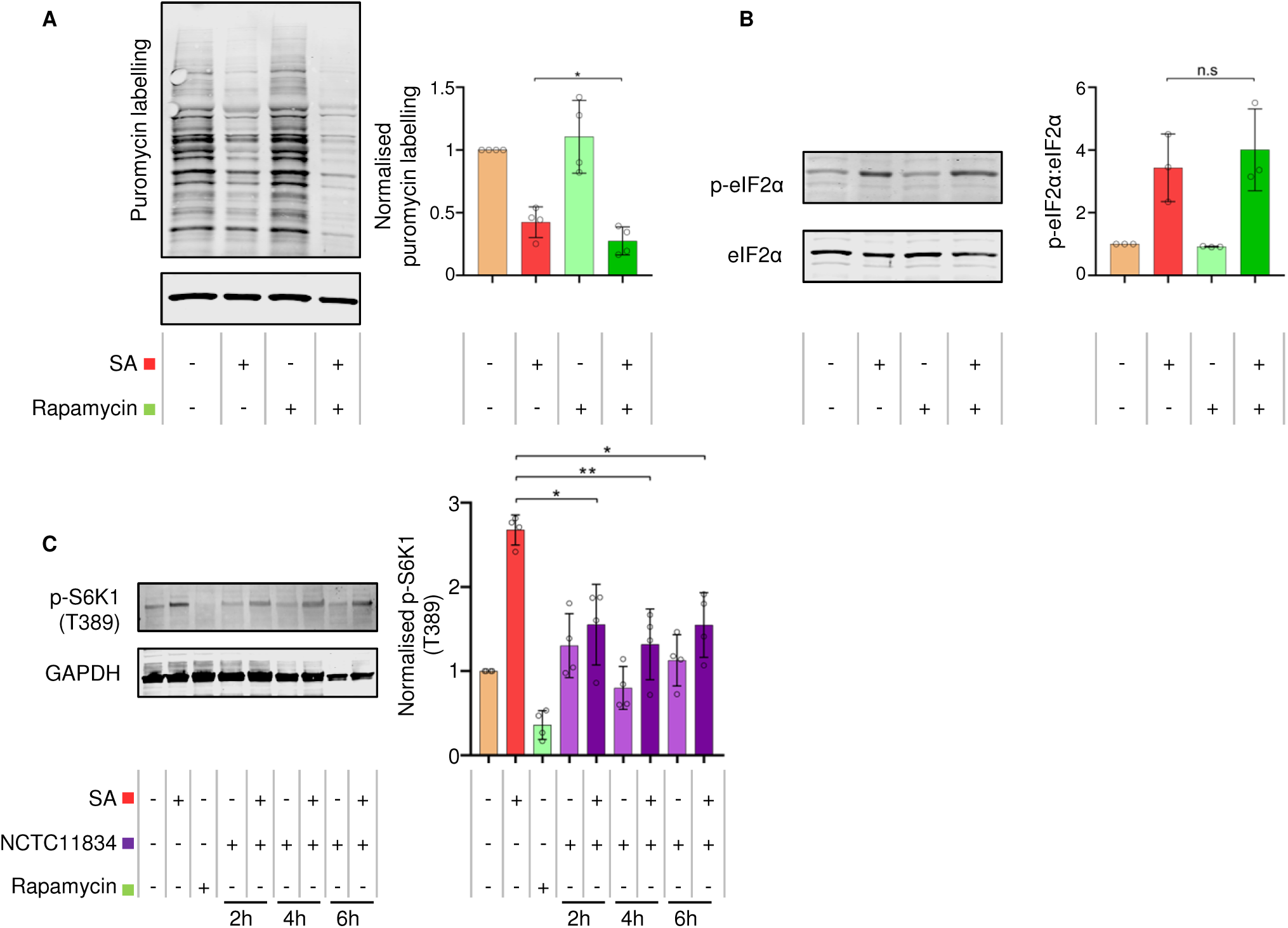
Rapamycin treatments exert the same effect on translation during oxidative stress as *P. gingivalis* and *P. gingivalis* attenuates stress-induced p-p70-S6-Kinase (T389). (A) Relative rate of protein synthesis as measured by puromycin uptake (left) and relative concentration compared to GAPDH in H357 cells treated with rapamycin and sodium arsenite as determined by immunoblotting (right). (B) Levels of phosphorylated eIF2α (left) and ratio of phosphorylated to total eIF2a as determined by immunoblotting (right) (mean ± SD, n=3). (C) Levels of p-p70-S6K1 (T389) (left) and p-p70-S6K1 (T389) concentration relative to GAPDH as determined by immunoblotting (right) (mean ± SD, n=4). **, *P* ≤ 0.01; *, *P* ≤ 0.05 according to Kruskal-Wallis with Conover-Inman *post-hoc*.

To observe the impact of mTOR degradation on translation inhibition during oxidative stress, downstream mTORC1 targets were investigated. Rapamycin resulted in a 0.64-fold decrease in the levels of phosphorylated p-p70-S6K1 (T389), whereas oxidative stress induced an increase of 1.67-fold. In contrast, whilst *P. gingivalis* infection alone did not result in altered levels of p-p70-S6K1 (T389), infection in the presence of oxidative stress caused a 0.4-fold decrease at all timepoints investigated (Fig 5C) suggesting that the phosphorylation activity of mTORC1 is downregulated by infection during stress.

### Secreted *P. gingivalis* proteases, gingipains, mediate heightened translational repression during stress

The findings thus far suggest that *P. gingivalis* can heighten translational repression during oxidative stress via a secretory factor. The impact of gingipains on translational control during oxidative stress and infection was therefore probed in the presence of the gingipain specific inhibitors TLCK (Lysine-specific, kgp) and Leupeptin (Arginine-specific, rgp). Both Leupeptin and TLCK either alone or in tandem, inhibited the ability of the conditioned media to heighten translational stalling during oxidative stress (Fig 6A).

**FIG 6.**
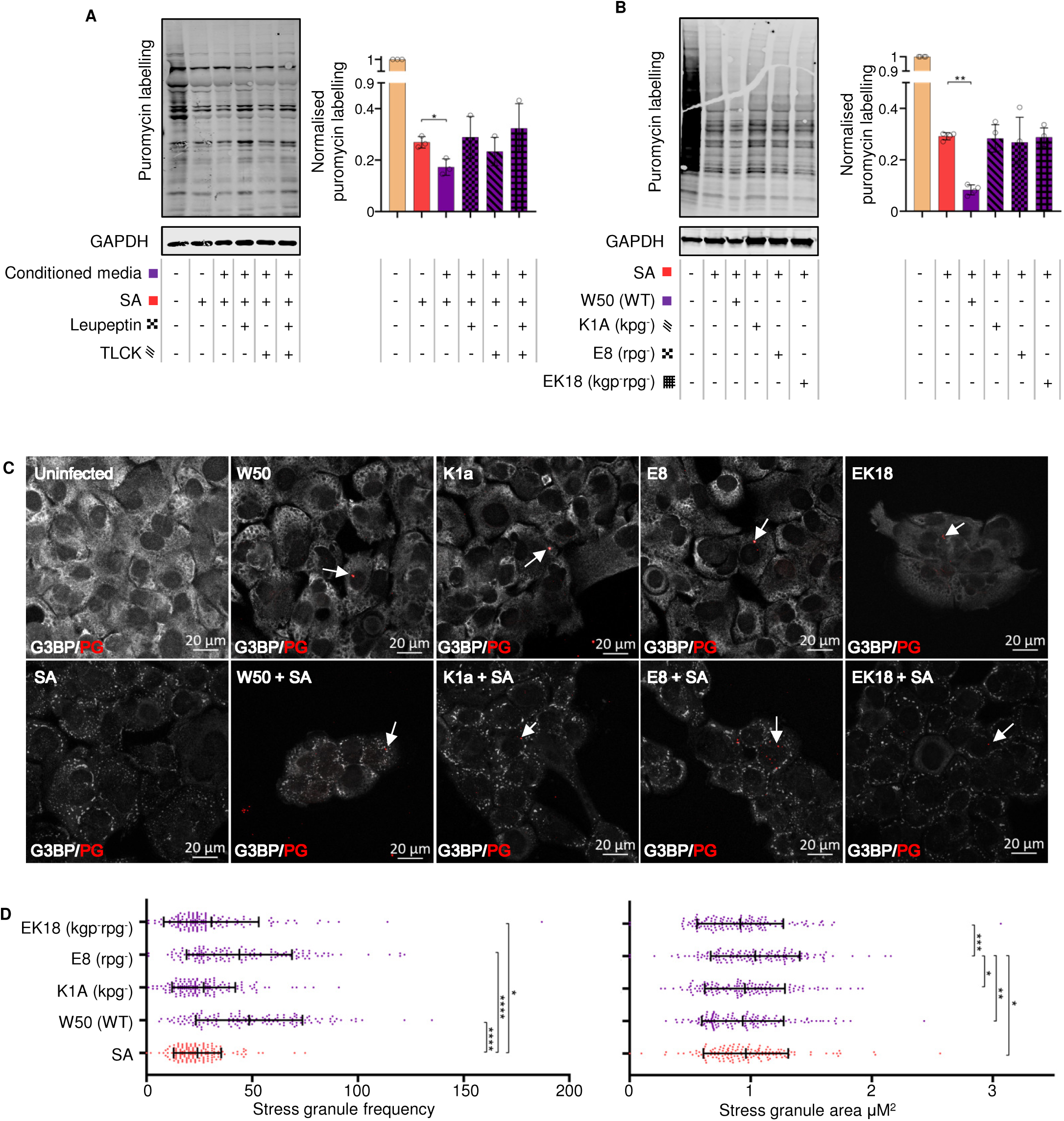
Secreted *P. gingivalis* proteases, gingipains, mediate heightened translational repression during infection and stress. (A) Relative rate of protein synthesis as determined by puromycin uptake (left) and concentration relative to GAPDH (right) when H357 cells were treated with *P. gingivalis* (NCTC11834) conditioned media in the presence or absence of leupeptin and TLCK and (mean ± SD, n=3) (B) with *P. gingivalis* (W50, K1A (*kgp^-^*), E8 (*rgp^-^*) and EK18 (*rgp^-^kgp^-^*); (mean ± SD, n=4). (C) Stress granule formation was assessed by visualisation of G3BP1 (white) and *P. gingivalis* (red) by confocal microscopy using Z-stacks. (D) Average area and frequency of SGs found in cells (n=3, 50 cells per biological replicate). **** *P* ≤ 0.001; ***, *P* ≤ 0.001; **, *P* ≤ 0.01; *, *P* ≤ 0.05 according to Kruskal-Wallis with Conover-Inman *post-hoc*.

To further confirm the role of gingipains in translational attenuation a series of isogenic gingipain null mutants in *P. gingivalis* strain W50 were studied. Neither the wild type W50 strain nor the K1A, E8 and EK18 mutants induced a change in protein synthesis during infection in the absence of oxidative stress (Fig S5). In the presence of oxidative stress and W50, puromycin incorporation decreased 0.72-fold, compared to the oxidative stress only treated control; however the mutants were unable to elicit this phenotype (Fig 6B).

These findings implicate gingipains in *P. gingivalis* mediated heightened translational repression during oxidative stress and hence the effect of gingipains on stress granules was investigated. Wild-type W50 strain induced an increase in stress granules during oxidative stress that was comparable to the NCTC11834 strain (Fig S6). Neither the wild-type nor the mutants induced stress granules or inhibited their formation during oxidative stress (Fig 6C). In oxidative stress treated cells, both the wild-type W50 and gingipain mutants E8 and EK18 induced an increase in stress granule frequency which was not observed in K1A infected cells. Neither the wild-type, K1A nor EK18 changed the average stress granule area, whereas surprisingly the E8 mutant increased the area of stress granules (Fig 6D). Taken together, these findings indicate that both lysine- and arginine-specific gingipains are accountable for *P*. *gingivalis* mediated heightened translation repression during oxidative stress, with the lysine-specific gingipain inducing the increased stress granule frequency.

### *P. gingivalis* infection dampens stress-induced tubulin acetylation

As changes to stress granule frequency and area were observed, the underlying mechanism of stress granule assembly was investigated. Assembly of mature stress granules requires aggregation of components into smaller foci along polymerising microtubules (34, 52). Here the integrity of the microtubule network was determined. Visualisation of α-tubulin showed no qualitative changes to the structure of α-tubulin following cell treatment with oxidative stress only or with *P. gingivalis* (NCTC11834; MOI 1:100, t=2h; with and without oxidative stress) when compared to the total lack of structure observed with the positive control nocodazole (Fig 7A).

**FIG 7.**
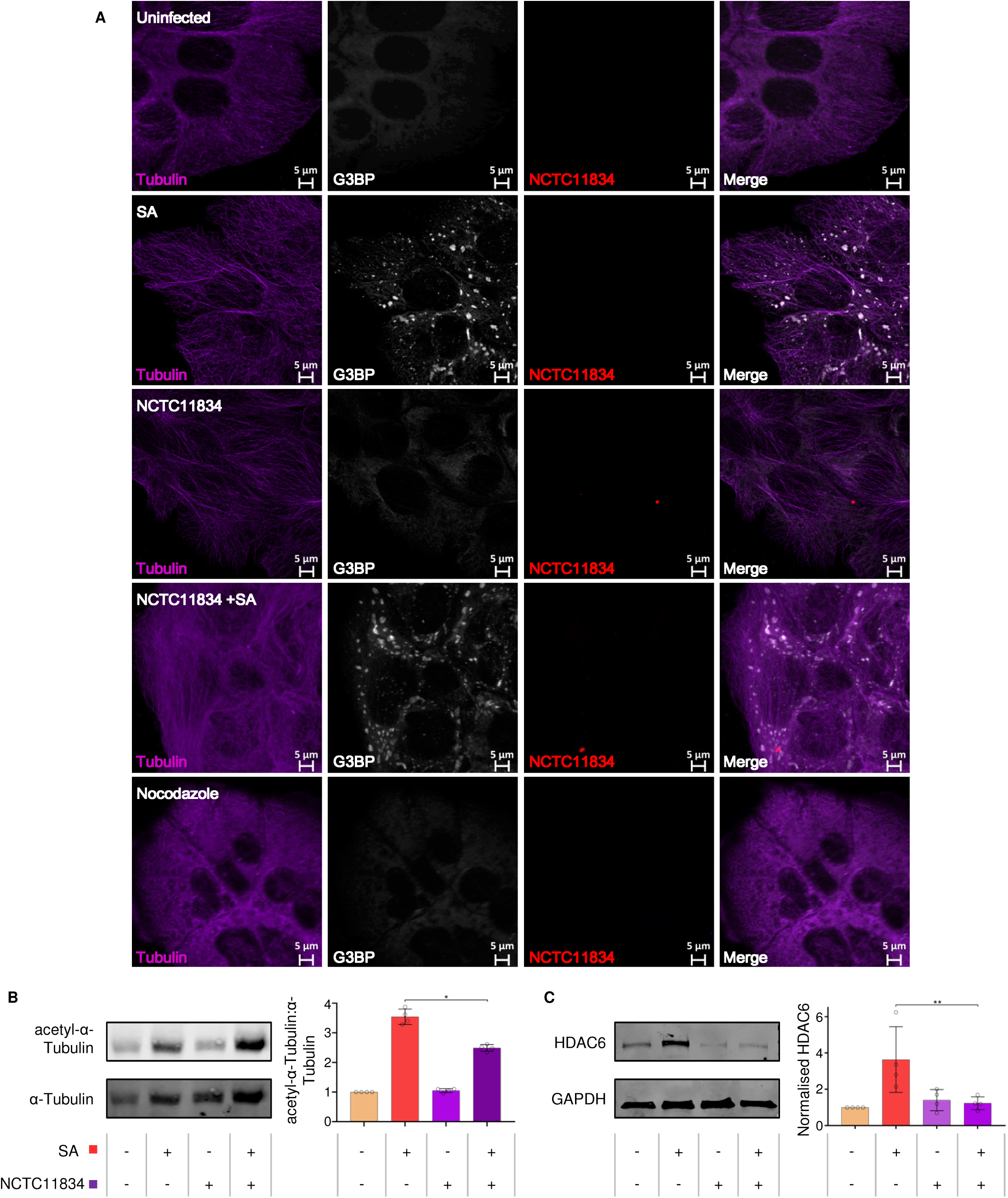
*P. gingivalis* infection dampens stress-induced tubulin acetylation. (A) Stress granule, a-tubulin integrity and *P. gingivalis* were visualised using confocal microscopy following challenge of H357 cells with *P. gingivalis* (NCTC11834) in the presence or absence of sodium arsenite. (B) Expression levels of a-tubulin (left) and ratio of acetyl-a-tubulin to a-tubulin (right; mean ± SD, n=4). (C) Expression of HDAC6 (left) and concentration relative to GAPDH (right; mean ± SD, n =3) as determined by immunoblotting. **, *P* ≤ 0.01; *, *P* ≤ 0.05 according to Kruskal-Wallis with Conover-Inman *post-hoc*.

As the function of tubulin can be modified post-translationally, the levels of acetyl-α-tubulin were monitored using immunoblotting. Both untreated cells and those infected with *P. gingivalis* (NCTC11834; MOI 1:100, t=2h) displayed basal level acetylation. Oxidative stress resulted in a 3.6 fold increase in acetylation; a response which was dampened (0.3-fold) when cells were infected with *P. gingivalis* prior to the addition of oxidative stress (Fig7B).

To investigate the means of tubulin deacetylation during oxidative stress, the expression of the principal tubulin deacetylation enzyme, HDAC6, was determined (Fig 7C). *P. gingivalis* infection (NCTC11834; t=2h) did not raise HDAC6 above basal levels whilst oxidative stress increased the levels of HDAC6 (3.6 fold). This phenotype was however not observed when infection was coupled with oxidative stress (Fig 7C) suggesting that the lowered tubulin acetylation observed during *P. gingivalis* and oxidative stress was independent of increased HDAC6 expression.

## DISCUSSION

In recent years, ISR signalling and translational control during stress have garnered increased interest within the remit of host immune responses. These pathways are capable of inducing a wide variety of outcomes at the cellular level, which subsequently feed into the organismal systemic responses (53). As such, these signalling cascades offer a promising target for pathogens to manipulate. Both bacteria and viruses have been shown to influence the ISR, thereby reprogramming a variety of host responses and enabling the generation of a favourable replicative niche (Reviewed in (20, 36)). This study aimed to investigate the crosstalk between host translational control during stress and *P. gingivalis* infection and the potential wider impact upon periodontal disease progression.

Previously, *P. gingivalis* infection has been shown to activate the UPR in human umbilical cord vein endothelial cells (47). Given that one arm of the UPR feeds into the ISR (48), we hypothesised that *P. gingivalis* infection may also activate the ISR. However, over a period of two, four, six and 24h, ISR activation was not observed, as evidenced by a lack of p-eIF2a or translational repression (Fig 1A and B); both core components of the active ISR (18). Furthermore, infection over the same time period did not result in the aggregation of G3BP into stress granules, a downstream marker of translational repression brought on by ISR activity (Fig 1C). Whilst it cannot be formally excluded that these responses might be cell type specific, with human umbilical cord vein endothelial cells previously used (47) in contrast to the squamous oral epithelial cell carcinoma cells used here, it is possible that the UPR may have been active independent of translational attenuation (discussed further in (54)).

Having established that *P. gingivalis* infection alone did not stimulate the ISR, the impact of infection coupled with oxidative stress was investigated, as host inflammatory responses are known to induce the production of reactive oxygen species following neutrophil activation (55). Sodium arsenite, one of the most well-characterised ISR activating stressors, induces oxidative stress via HRI kinase (56). In this study, the high levels of inflammation characteristic of periodontitis and caused by *P. gingivalis* infection (57) coupled with the expression of oxidative stress resistance genes by *P. gingivalis* (58) made sodium arsenite an attractive and relevant stress.

Previous studies (15) and analysis of *P. gingivalis* treated cells here showed that 20% of cells of a population are invaded between two and four hours post infection (Fig 1E). In the presence of oxidative stress, bacterial invasion was found to increase 2-fold. Although the exact cause of this increase remains to be elucidated, *P. gingivalis* is known to express its own oxidative stress resistance genes (58) and to actively protect host cells against reactive oxygen species via the host antioxidant glutathione response (59). *P. gingivalis* is further protected by a layer of hemin on its cell surface which acts as a buffer against oxidative radicals and increases *P. gingivalis’* resistance to host oxidative stress (60, 61). *P. gingivalis* can utilise a multitude of defences against oxidative stress whilst simultaneously upregulating host antioxidant pathways. This coupled with the fact that sodium arsenite exposure can decrease mammalian membrane integrity (62) may underpin the increased invasion observed during sodium arsenite induced oxidative stress.

Oxidative stress, as expected, resulted in translational repression (56). *P. gingivalis*, in the presence of oxidative stress, exacerbated translational repression (Fig 1D) and increased stress granule frequency (Fig 2B). Previous studies looking at *S. flexneri* infection, have implicated mTORC1 inhibition due to the membrane damage caused by bacterial internalisation, in stress granule modulation and translational dysregulation (38, 45). An increase in stress granule frequency has also been reported during chemical mTOR inhibition (63). The research from these groups’ findings suggests that mTORC1 has a role in increased translational attenuation and stress granule frequency and is supported by the involvement of mTORC1 as a key regulator of translation (64), with its inhibition leading to polysome disassembly and subsequent translational stalling (65). Although *P. gingivalis* can inhibit and degrade mTOR (15, 46), *P. gingivalis* alone did not lead to translational attenuation. A similar result was observed during rapamycin treatment. These differences in translational attenuation could reflect the variable outcomes of mTORC1 inhibition under different conditions (66). The effects of *P. gingivalis* mediated inhibition and degradation on translation may therefore only become apparent in presence of another stress as seen here where *P. gingivalis* heightened oxidative stress-induced translational attenuation.

Stress granules are formed by sequestration of stalled mRNPs into smaller foci, which in due course fuse into larger aggregates (52). The increased frequency of stress granules observed in this study may be due to *P. gingivalis* dysregulating stress granule aggregation and partially excluding eIF3b from the stress granules (Fig 2C and E). This is corroborated by reports that *S. flexneri* can selectively cause delocalisation of eIF3b from stress granules during exogenous stress, in a manner dependent on mTORC1 inactivation (45). The movement of eIF3b is regulated by mTORC1, which phosphorylates S6K1 at T389, releasing S6K1 from eIF3b (67). Oxidative stress-induced p-S6K1 (T389) (68) was decreased by *P. gingivalis* infection (Fig 5C), probably owing to inhibition or degradation of mTORC1 (15, 46). Therefore, decreased p-S6K1 (T389) could account for the lack of eIF3b in stress granules during oxidative stress and *P. gingivalis* infection (Fig 2E) and further supports the role of mTORC1 in the exclusion of eIF3b from stress granules.

Aggregation of stress granules requires constant retrograde transport along functioning microtubules (46, 52). Nocodazole, a chemical which disrupts microtubule assembly (69), increases the frequency of stress granules (70). *P. gingivalis* is known to degrade cytoskeletal protein components such as β-actin (15, 71). Therefore, we next determined if the increase in frequency of stress granules observed following *P. gingivalis* infection in the presence of oxidative stress was the result of tubulin degradation. Infection did not result in visible changes to the microtubular network compared to nocodazole treated cells (Fig 7A). However, microtubule network activity can also be controlled via post-translational modifications such as acetylation and phosphorylation (72), with hyper-acetylation of the cellular tubulin network at lysine 40 of α-tubulin reported during stress (14). α-tubulin hyper-acetylation stimulates increased binding and activity of the microtubule motor proteins dynein and kinesin, involved in the movement of stress granules (73–75). Here *P. gingivalis* lowered the levels of α-tubulin acetylation during infection and oxidative stress (Fig 7B), which was independent of increased expression of HDAC6, the major α-tubulin deacetylase (76). Furthermore, HDAC6 is a critical stress granule component, ablation of which inhibits stress granule assembly (77). Hence the decreased tubulin acetylation and lowered HDAC6 expression may be influencing the modulated stress granule formation seen here during *P. gingivalis* infection and oxidative stress.

During ISR activation, translational attenuation, due to a range of stressors, is mediated by the phosphorylation of eIF2α (18). Given the fundamental role of p-eIF2α in translational control during oxidative stress, dysregulating eIF2α phosphorylation is a mechanism by which many viruses hijack host translational function (Reviewed in (78)). In this study, despite infection by *P. gingivalis* heightening translational repression during oxidative stress, no increase in p-eIF2α was observed (Fig 3A). These results were corroborated by the inability of ISRIB to rescue translational function and to inhibit stress granule assembly during oxidative stress and infection (Fig 3B/C). ISRIB functions to induce a conformational change in eIF2B, antagonising the inhibitory effects of p-eIF2α (79, 80). Hence the data point towards a mechanism independent of eIF2α as the mediator of the heightened translational repression seen during *P. gingivalis* infection and oxidative stress.

As the heightened translational repression was eIF2α independent and downstream mTORC1 targets were altered, rapamycin, a potent mTOR inhibitor, was used to further probe the pathway. During oxidative stress, rapamycin induced the same phenotype as *P. gingivalis* infection, heightened oxidative stress-induced translational stalling independently of p-eIF2α (Fig 5A/B), further supporting the contributory role for mTORC1. *P. gingivalis* gingipains which have been shown to both degrade and inhibit mTOR (15, 46), are expressed as cell surface anchored proteins or in the secretome of *P. gingivalis*, where they exist both freely or packaged within OMVs (13, 81, 82). Given that *P. gingivalis* conditioned media and OMVs exhibited a similar phenotype to internalised bacteria, the role of gingipains was next investigated. Inhibition of gingipains in conditioned media by gingipain-specific inhibitors inhibited the heightened translational attenuation observed with the conditioned media (Fig 6A). This inability to induce further translational repression was also observed using gingipain-knockout mutants (Fig 6B). These findings implicate both the arginine- and lysine-specific gingipains in an extra and intracellular manner and is possibly due to mTORC1 inhibition via the PI3K pathway as reported by Nakayama and colleagues (46). When the impact of these gingipain-knockout mutants on stress granule formation was investigated, the lysine gingipain-knockout (kgp) failed to increase stress granule frequency (Fig 6D) which could reflect the requirement of intracellular *P. gingivalis* secreted lysine-specific gingipains for mTOR degradation (15). The requirement for internalisation to enact the function of stress granule modulation may account for the less marked increase in stress granule frequency in the *P. gingivalis* negative cells of the exposed population, as OMVs containing gingipains only enter around 8% of cells (83). In contrast 40% of cells were infected in this study, when exposed to *P. gingivalis* and oxidative stress (Fig 2B). Furthermore, as gingipains are secreted following *P*. *gingivalis* invasion and internalisation of host cells (83, 84), it could increase the concentration of intracellular lysine-specific gingipain, compared to conditioned media and OMV treatment. Taken together these findings suggest that while both gingipains can heighten translational repression, the lysine-specific gingipain is the main effector of stress granule modulation and works most efficiently after invasion.

This study has for the first time demonstrated that the periodontopathogen *P. gingivalis* dysregulates translational control and stress granule formation during oxidative stress, a condition phenotypic of the chronic inflammatory environment induced during periodontitis and caused by *P. gingivalis* (3) (Illustrated in Fig. 8). These findings suggest a novel pathogenic mechanism employed by *P. gingivalis* to modulate host response and given that these pathways feed into cellular survival and the wider immune, and inflammatory response (53), can contribute to the immune subversive nature of *P. gingivalis.* Furthermore, dysregulation of the ISR, translational control and stress granule dynamics have been implicated in a range of diseases from cancer to neurodegeneration (53, 85, 86). Further investigations building on the data presented here may therefore provide insight into the relationship between systemic *P. gingivalis* infection and other diseases.

**FIG 8.**
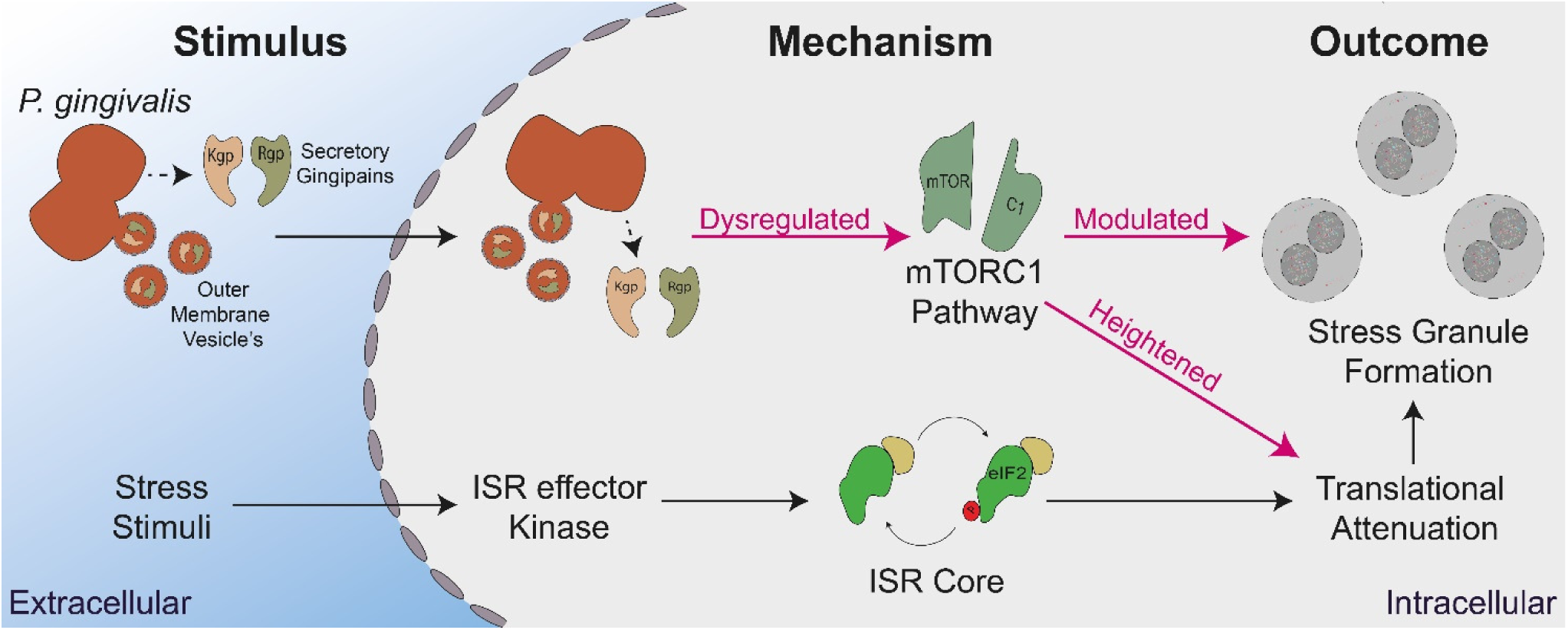
Summary of *P. gingivalis* interactions with host stress-induced translational control. Stress stimuli, such as oxidative stress, activate ISR effector kinases which phosphorylate eIF2a, resulting in translational attenuation and stress granule formation. *P. gingivalis* secretes gingipains and outer membrane vesicles in a both extra and intracellular manner, which dysregulate the mTORC1 pathway leading to heightened translational attenuation and modulated stress granule formation.

## MATERIALS AND METHODS

### Reagents

All cell culture reagents unless otherwise stated were from Sigma/Merck Life Science UK LTD (Dorset, UK).

### Cell culture

The oral squamous carcinoma derived cell line (H357) was maintained in Dulbecco’s modified Eagle’s medium (DMEM; Gibco, Fisher Scientific, Loughborough UK) supplemented with 10% foetal bovine serum (FBS) and 2 mM L-glutamine (Glu) in a humidified environment (5% CO_2_, 37°C). Cells were passaged when ∼75% confluent by trypsinization and cell viability were assessed using trypan blue exclusion method as previously described (15).

### Bacterial strains and culture

Bacterial strains used in this study include *P. gingivalis* NCTC11834, W50 (ACTC 53978) and the derivative W50 isogenic mutants K1A (*kgp*::*Em*), E8 (*rgpA::Em rgpB::Tet*) (87) and EK18 (*rgpA::Em rpgB::Tet kgp::Chlor*) (15). All strains used were a kind gift from Professor G. Stafford (School of Clinical Dentistry, University of Sheffield, UK).

*P. gingivalis* were grown and maintained on fastidious anaerobe agar (Lab M, Bury, UK) containing oxylated horse blood (5%(v/v); TCS Biosciences, Buckingham, UK) and supplemented with antibiotics as required under anaerobic conditions (10% CO_2_, 10% H_2_ and 80% N_2_) at 37°C. Bacteria were subcultured every 3-4 days for maintenance. Throughout this study, bacteria were used to infect cells when no older than 3-4 days old post-subculturing. *P. gingivalis* were grown as liquid cultures in brain heart infusion broth (BHI, Difco laboratories, East Molesey, Surrey, UK) supplemented with 0.5%(w/v) yeast extract, hemin (5µg/ml), vitamin K (0.5µg/ml) and cysteine (0.1%)(w/v). Purity of liquid cultures was confirmed by Gram staining before use.

### Bacterial infection, oxidative stress induction and cell treatments

H357 were seeded at a density of 6×10^4^ cells/cm^2^ on coverslips or at 3.6×10^4^ per cm^2^ in tissue culture flasks in DMEM/Glu/FBS, following which cells were incubated (5% CO_2_ 37°C) and allowed to adhere overnight. After replacement of overnight media with fresh media, cells were challenged with *P. gingivalis* at a multiplicity of infection (MOI) 1:100 at the time points as detailed below. Oxidative stress was induced using sodium arsenite (SA, 250µM) which was added for the final 30 min of infection. Cells were also treated with or without ISRIB (200nM, 30 min), Nocodazole (200nM, 30 min), Rapamycin (400nM, t=1h) or *P. gingivalis* (NCTC11834) derived Lipopolysaccharide (LPS) (at 1, 5 or 10µg/mL, t=2h). Uninfected cells were included as control.

After treatment, for Western blotting, adherent cells were washed with phosphate buffered saline (PBS), before the addition of lysis buffer (PBS supplemented with 10 %(v/v) PhosStop (Roche), 10 %(v/v) complete EDTA-free protease inhibitors and 0.1 %(v/v) SDS). Total proteins were extracted using a cell scraper and cell lysates were stored at -80°C for a minimum of one hour, or overnight after which proteins were recovered by centrifugation (17 200xg, 14 min, 4°C) and stored at -80°C until required. Total protein extracts were quantified using the Qubit^TM^ protein assay (ThermoFisher) according to manufacturer instructions and expression levels of proteins of interest probed by Western blotting. For immunofluorescence analysis, cells were fixed as detailed below.

### Isolation of *P. gingivalis* outer membrane vesicles (OMVs)

*P. gingivalis* OMVs were extracted as previously described (88). *P. gingivalis* were grown to late exponential phase overnight in liquid culture as outlined above. The next day, cultures were adjusted to OD_600_ of 1.0 following which they were subjected to centrifugation (8,000xg, 5 min, 4°C). The resulting supernatant was filtered-sterilised (0.22µM) and centrifuged (100,000xg, 2h, 4°C), after which the supernatant was discarded, and the pellet resuspended in PBS. Protein content was determined as outlined above and the resulting quantified OMVs used to challenge H357 cells.

### Generation of *P. gingivalis* conditioned media and gingipain inhibition

To determine the effect of *P. gingivalis* secreted components, H357 cells were infected (MOI 1:100) as described above after which the conditioned media was recovered and filtered (0.22µM) to remove bacteria and other particulate matter. Untreated adherent H357 cells were then challenged with the recovered conditioned media for 2h. For gingipain inhibition studies, oral epithelial cells were challenged with conditioned media supplemented with either leupeptin (0.2mM) or Na-Tosyl-Lysine Chloromethyl Ketone (TLCK, 0.5mM) after which total protein was extracted and levels of proteins of interest were probed by Western blotting.

### Western blotting

For western blotting, total protein extracts were subjected to SDS page electrophoresis using 4-20% polyacrylamide gradient gels (Bio-Rad, Watford, UK) and transferred to nitrocellulose membranes using a Trans-blot Turbo transfer system (Bio-Rad). Membranes were blocked in either (5% w/v) bovine serum albumin (BSA) or powdered milk prepared in Tris Buffered Saline (TBS; 37mM NaCl, 20mM Tris, pH 7.6) supplemented with 0.1%(v/v) Tween 20 (TBST) for 1 hour at room temperature before incubation with primary antibodies overnight at 4°C. Primary antibodies used include: puromycin (1:500; clone 12D10, MABE343, Merck), phosphorylated eIF2a (serine 51) (1:500, 44-728G, Invitrogen, Fisher Scientific), eIF2a (1:500, ab181467, Abcam), G3BP (1:500, ab56574, Abcam), eIF3b (1:500, ab133601, Abcam), phosphorylated p70-S6 Kinase (Threonine 389) (1:200, 108D2, Cell Signalling), phosphorylated 4E-BP1 (Threonine 37/46) (1:200, 236B4, Cell Signalling), a-tubulin (1:500, 2144, Cell Signalling), acetyl-a-tubulin (1:500, 1215, Cell Signalling) GAPDH (1:10,000, G9545, Invitrogen, Fisher Scientific) and GAPDH (1:10,000, PL0125, Invitrogen, Fisher Scientific). After washing with TBST (3 x 5 min), membranes were incubated with the corresponding fluorescent conjugated secondary antibodies for 1h (1:10,000, Li-Cor, location, UK). Proteins were visualised using a Li-Cor Odyssey infrared imager (Li-Cor) and quantified using Image Studio Lite software (Li-Cor).

### Puromycin incorporation assay

The relative rates of protein synthesis were determined using the non-radioactive fluorescence activated surface sensing of translation assay as described previously (89). Briefly, post-treatment cells were incubated in culture media containing puromycin (91µM) and emetine (208µM) for 5 minutes (5% CO_2_, 37°C). Cells were then washed twice with PBS containing cycloheximide (355µM) and total protein was extracted as detailed above following which puromycin uptake was probed by western blotting.

### Immunocytochemistry

Methanol-fixed cells were washed with PBS supplemented with Tween 20 (0.5% v/v; PBST), following which the cells were blocked in PBS supplemented with BSA (1% w/v) for a minimum of 1 hour at room temperature before incubation with primary antibodies overnight. The following primary antibodies were used: G3BP (1:500, ab56574, Abcam), eIF3b (1:500, ab133601, Abcam), a-tubulin (1:500, ab6161, abcam) and *P. gingivalis* (1:500, a kind gift from Prof. G. Stafford, University of Sheffield Dental School). After washing with PBST (3 x 5 min), membranes were incubated with corresponding fluorescent Alexa fluor^TM^ conjugated secondary antibodies for one hour at room temperature. Cells were washed with PBST and mounted using ProLong Gold^TM^ antifade mountant containing DAPI (ThermoFisher). Protein localisation was visualised using a Zeiss LSM800 microscope (Carl Zeiss, Cambridge, UK). Images were captured using ZenBlue software, either a 40x or 63x plan-apochromat oil objective and a laser with maximum output of 10mW at 0.2% laser transmission. Stress granule frequencies, area and co-localization were quantified using the analysis module of Zeiss ZenBlue software (Carl Zeiss).

### Statistical Analysis

Significance between groups was analysed using the StatsDirect software package (Statsdirect Ltd, Birkenhead, UK). Data was first subjected to a Shapiro-Wilks test where data was considered parametric if p<0.05. All data was found to be non-parametric. Significance between unpaired groups was determined using a Kruskal-Wallis test, which if significant was followed by a Conover-Inman *post-hoc* test. Significance was set at p≤ 0.05; **** *P* ≤ 0.001; ***, *P* ≤ 0.001; **, *P* ≤ 0.01; *, *P* ≤ 0.05.

## Declaration of competing interest

The authors declare that they have no competing interest.

## Acknowledgements

The authors would like to thank Professor Tom Smith and Dr Rachel Hodgson for fruitful discussions and Professor Graham Stafford for the kind gift of the *P. gingivalis* gingipain null mutant strains. The authors would also like to gratefully acknowledge the Biomolecular Sciences Research Centre and Sheffield Hallam University, Sheffield, UK for funding this work.

## Author Contributions

AK, SC, NC and PS conceived and designed the experimental plan. AK carried out the laboratory work and data analysis. AK, SC, NC, PS analysed the data, wrote, edited and revised the manuscript. All authors approved the final manuscript.

---

**FIG S1.**
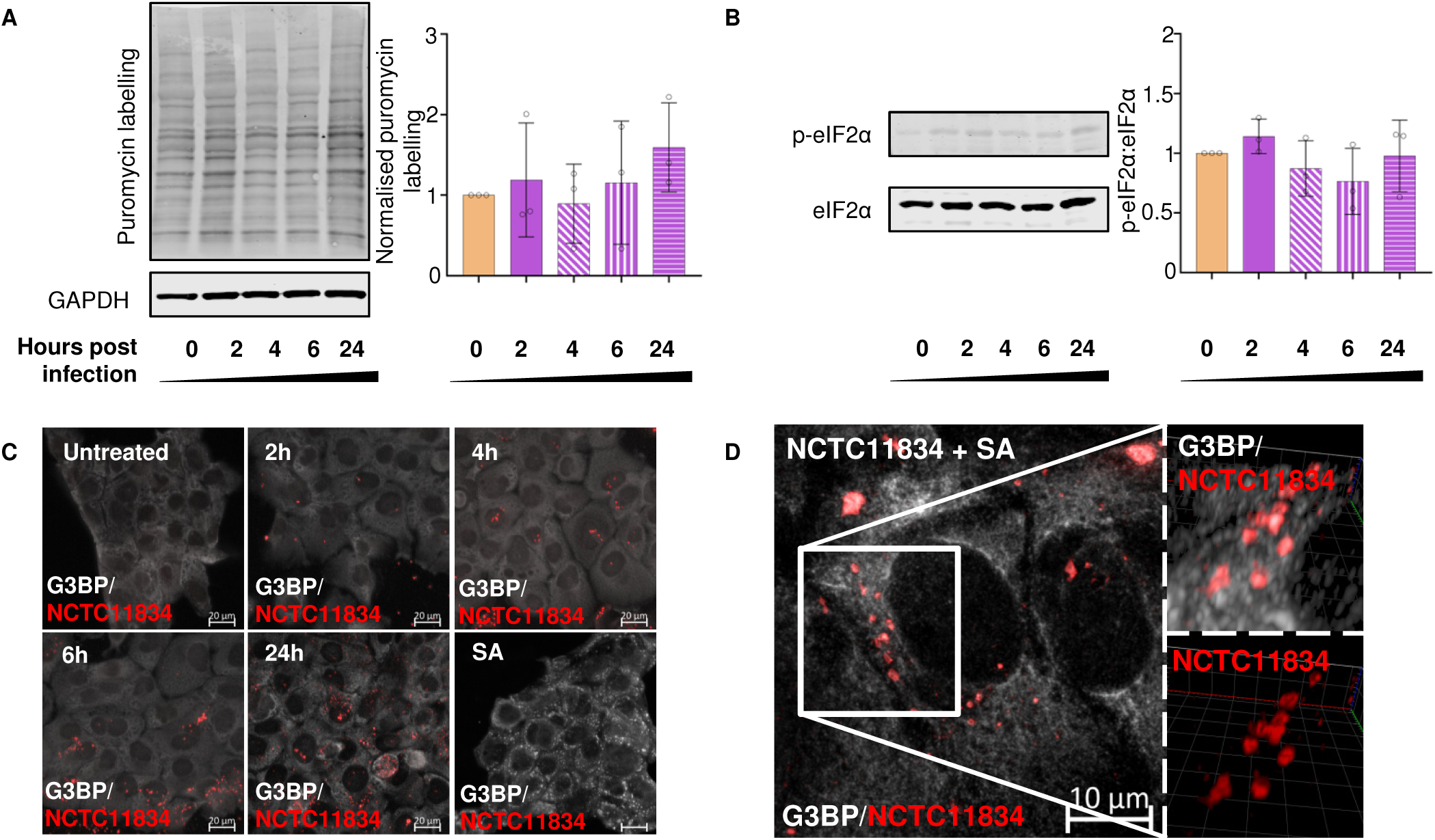
*P. gingivalis* does not induce ISR activation. (A) H357 cells were left untreated, infected with *P. gingivalis* (NCTC11834, MOI 1:100, t=2h to 6h) in the presence or absence of sodium arsenite as shown. Relative rate of protein synthesis as measured by puromycin uptake (left) and concentration relative to GAPDH (right) (B) Levels of phosphorylated eIF2α (p-eIF2α) were probed using immunoblotting (left) and the ratio to phosphorylated to total eIF2a was determined (right). GAPDH was included as a loading control (mean ± SD, n=3). (C) Stress granule formation was assessed by visualisation of G3BP (white) and *P. gingivalis* (red). (D) H357 cells were infected with *P. gingivalis* (NCTC11834, MOI 1:100, t=24h). G3BP (white) and *P. gingivalis* (red) were visualised using immunofluorescence confocal microscopy. No significant differences in means were found with a Kruskal-Wallis test.

**FIG S2.**
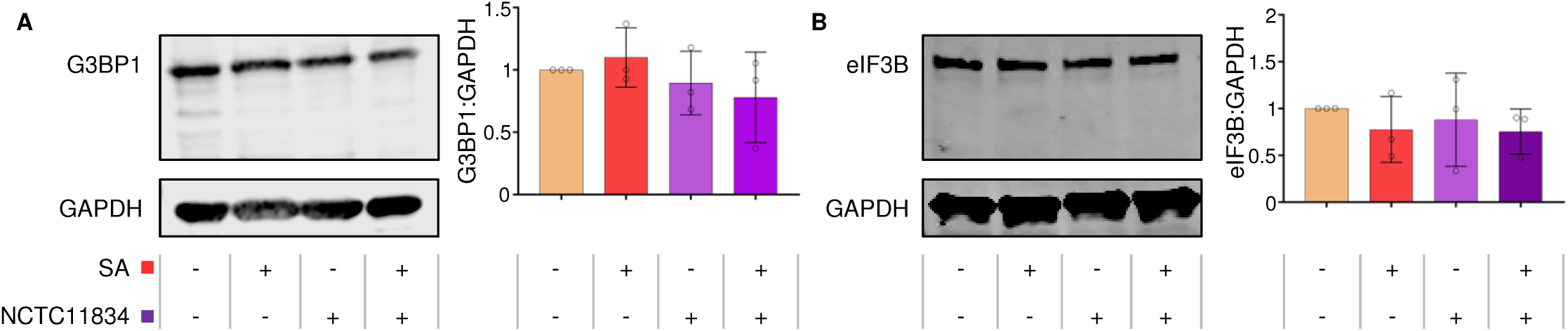
*P. gingivalis* and exogenous stress do not alter G3BP or eIF3B expression. H357 cells were left untreated or infected with *P. gingivalis* (NCTC11834, MOI 1:100, t=2h to 6h). Expression levels of (A) G3BP1 and (B) eIF3B were probed using immunoblotting. Concentration relative to the loading control GAPDH was first determined before being normalised to the untreated sample. Data are expressed as mean ± SD, n=3. No significant differences in means were found with a Kruskal-Wallis test.

**FIG S3.**
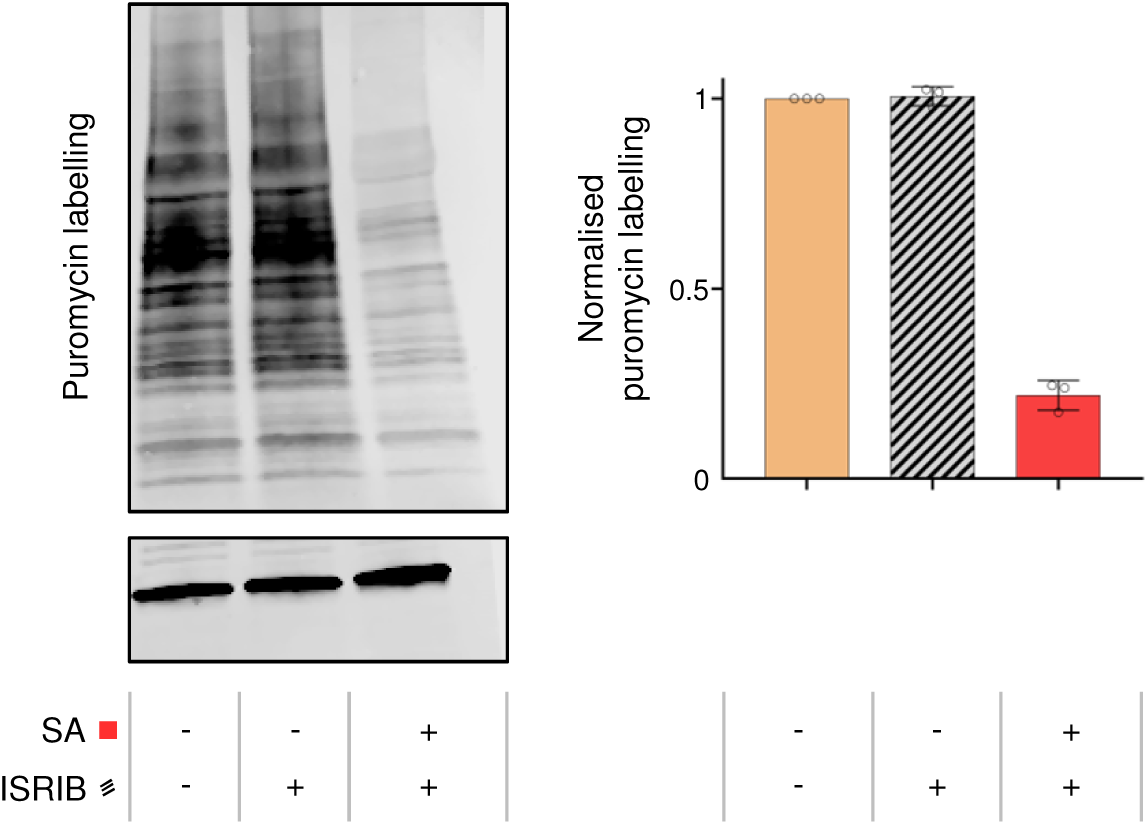
ISRIB treatment does not alter translation in H357 cells. H357 cells were treated with either ISRIB or sodium arsenite, following which the relative rate of protein synthesis, normalised first to the loading control GAPDH and then to control sample, was determined by immunoblotting for puromycin incorporation (mean ± SD, n=3). No significant differences in means were found with a Kruskal-Wallis test.

**FIG S4.**
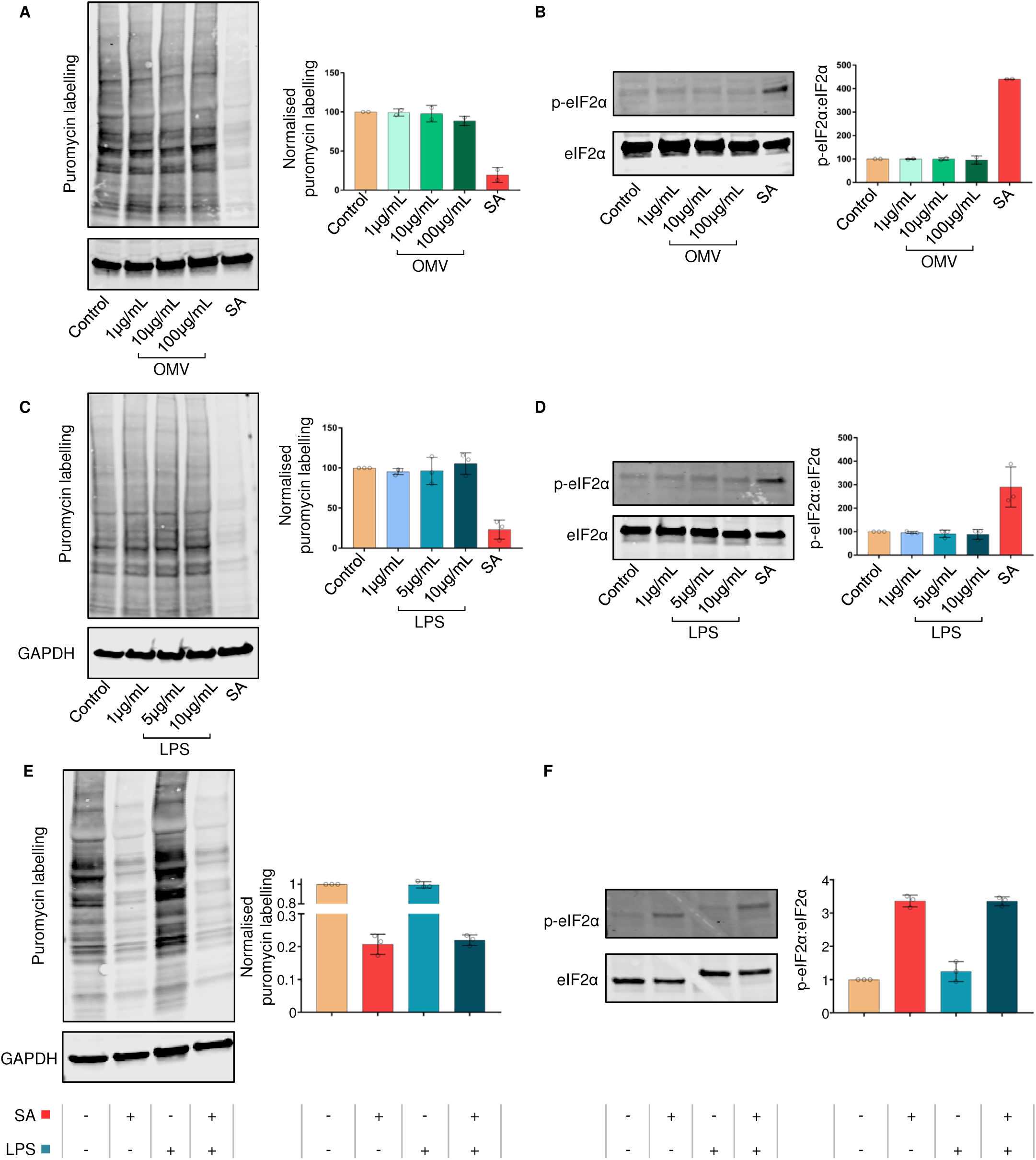
*P. gingivalis* outer membrane vesicles and lipopolysaccharide do not induce ISR activation. H357 cells were challenged with purified *P. gingivalis* OMV vesicles (1, 10 and 100µg/mL, t= 2h). Sodium arsenite was included as a positive control for ISR activation. (A) The relative rate of protein synthesis as measured by puromycin uptake and (B) the levels of phosphorylated eIF2α were probed using immunoblotting. To the right of each panel is a column graph of the quantified blot signals (mean ± SD, n=2). (C) H357 cells were challenged with purified *P. gingivalis* LPS (1, 5 and 10 µg/mL, t=2h). Sodium arsenite was included as a positive control for ISR activation. Following which the relative rate of protein synthesis measured by puromycin uptake and (D) the levels of phosphorylated eIF2α were probed using immunoblotting. To the right of each panel is a column graph of the quantified blot signals (mean ± SD, n=3). GAPDH was included as a loading control. (E) H357 cells were challenged with purified *P. gingivalis* LPS (10 µg/mL) with or without sodium arsenite for the final 30 min and the relative rate of protein synthesis measured by puromycin uptake (left) and concentration relative to GAPDH (right). (F) The levels of phosphorylated eIF2α (left) and the concentration of phosphorylated to total eIF2α (right) with LPS were probed using immunoblotting (mean ± SD, n=3). No significant differences in means were found with a Kruskal-Wallis test.

**FIG S5.**
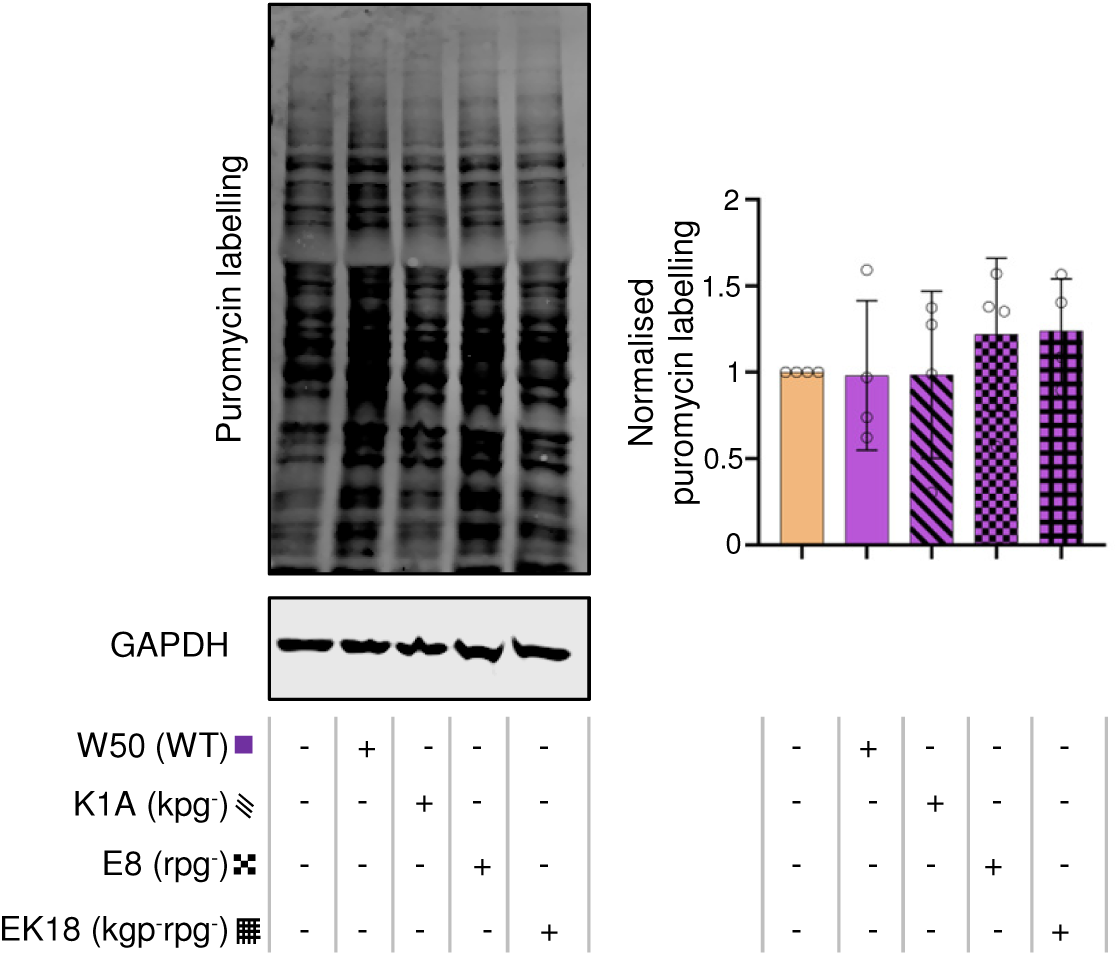
*P. gingivalis null* mutants do not alter protein synthesis. H357 cells were left untreated or infected with *P. gingivalis* (W50, K1A (*kgp^-^*), E8 (*rgp^-^*) and EK18 (*rgp^-^kgp^-^*), MOI 1:100, t =2h). Relative rate of protein synthesis was measured by immunoblotting for puromycin uptake. GAPDH was included as a loading control (mean ± SD, n=3). No significant differences in means were found with a Kruskal-Wallis test.

**FIG S6.**
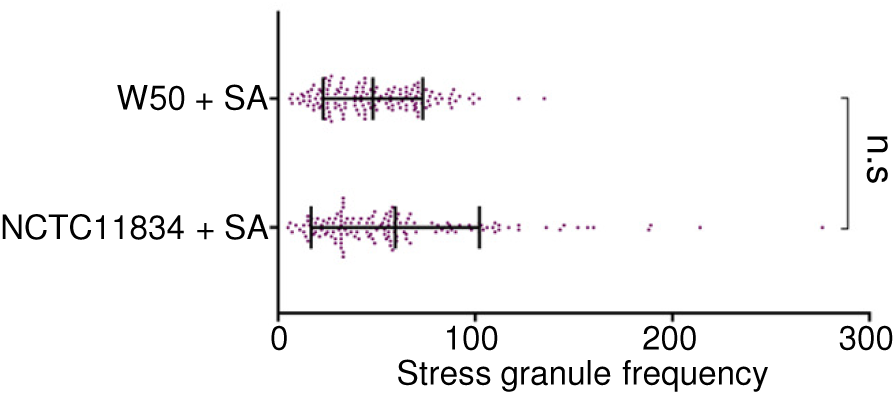
Comparison of stress granule frequency between NCTC11834 and W50 during stress. H357 cells were left untreated or infected by *P. gingivalis* (strains NCTC11834 and W50, MOI 1:100, t=2h) and treated with or without sodium arsenite for the final 30 min. Stress granule formation was assessed by visualisation of G3BP1 (white) and *P. gingivalis* (red) by confocal microscopy using Z-stacks. (n=3, 50 cells per biological replicate). No significant differences in means were found with a Kruskal-Wallis test.

## References

1. Marsh, PD. 2004. Dental Plaque as a Microbial Biofilm. Caries Res. 38:204–211. doi: 10.1159/000077756.

2. Socransky, SS, Haffajee, AD, Cugini, MA, Smith, C, Kent Jr, R,L. 1998. Microbial complexes in subgingival plaque. J Clin Periodontol 25:134–144. doi: 10.1111/j.1600-051X.1998.tb02419.x.

3. Pihlstrom, BL, Michalowicz, BS, Johnson, NW. 2005. Periodontal diseases. Lancet. 366:1809–1820. doi: 10.1016/S0140-6736(05)67728-8.

4. Tonetti, MS, Jepsen, S, Jin, L, Otomo-Corgel, J. 2017. Impact of the global burden of periodontal diseases on health, nutrition and wellbeing of mankind: A call for global action. J Clin Periodontol 44:456–462. doi: 10.1111/jcpe.12732.

5. Kebschull, M, Demmer, RT, Papapanou, PN. 2010. “Gum Bug, Leave My Heart Alone!”—Epidemiologic and Mechanistic Evidence Linking Periodontal Infections and Atherosclerosis. J. Dent. Res. 89:879–902. doi: 10.1177/0022034510375281.

6. Bingham, O, C., Moni, O, M. 2013. Periodontal disease and rheumatoid arthritis: the evidence accumulates for complex pathobiologic interactions. Curr Opin Rheumatol. 25:345–353. doi: 10.1097/BOR.0b013e32835fb8ec.

7. Preshaw, P, Alba, A, Herrera, D, Jepsen, S, Konstantinidis, A, Makrilakis, K, Taylor, R. 2012. Periodontitis and diabetes: a two-way relationship. Diabetologia. 55:21–31. doi: 10.1007/s00125-011-2342-y.

8. Gnanasekaran, J, Binder Gallimidi, A, Saba, E, Pandi, K, Eli Berchoer, L, Hermano, E, Angabo, S, Makkawi, H, Khashan, A, Daoud, A, Elkin, M, Nussbaum, G. 2020. Intracellular Porphyromonas gingivalis Promotes the Tumorigenic Behavior of Pancreatic Carcinoma Cells. Cancers. 12:2331. doi: 10.3390/cancers12082331.

9. Dominy, SS, Lynch, C, Ermini, F, Benedyk, M, Marczyk, A, Konradi, A, Nguyen, M, Haditsch, U, Raha, D, Griffin, C, Holsinger, LJ, Arastu-Kapur, S, Kaba, S, Lee, A, Ryder, MI, Potempa, B, Mydel, P, Hellvard, A, Adamowicz, K, Hasturk, H, Walker, GD, Reynolds, EC, Faull, RLM, Curtis, MA, Dragunow, M, Potempa, J. 2019. Porphyromonas gingivalis in Alzheimer’s disease brains: Evidence for disease causation and treatment with small-molecule inhibitors. Science Advances; Sci Adv. 5:eaau3333. doi: 10.1126/sciadv.aau3333.

10. Adams, Bü, Nunes, JM, Page, MJ, Roberts, T, Carr, J, Nell, TA, Kell, DB, Pretorius, E. 2019. Parkinson’s Disease: A Systemic Inflammatory Disease Accompanied by Bacterial Inflammagens. Frontiers in Aging Neuroscience; Front Aging Neurosci. 11:210. doi: 10.3389/fnagi.2019.00210.

11. Hajishengallis, G, Darveau, RP, Curtis, MA. 2012. The keystone-pathogen hypothesis. Nature Reviews Microbiology. 10:717. doi: 10.1038/nrmicro2873.

12. Hajishengallis, G, Lambris, JD. 2012. Complement and dysbiosis in periodontal disease. Immunobiology. 217:1111–1116. doi: 10.1016/j.imbio.2012.07.007.

13. Grenier, D, Choa, G, McBride, BC. 1989. Characterization of sodium dodecyl sulfate-stable Bacteroides gingivalis proteases by polyacrylamide gel electrophoresis. Infect Immun. 57:95–99. doi: 10.1128/IAI.57.1.95-99.1989.

14. Li, N, Collyer, CA. 2011. Gingipains from Porphyromonas gingivalis - Complex domain structures confer diverse functions. European Journal of Microbiology & Immunology. 1:41. doi: 10.1556/EuJMI.1.2011.1.7.

15. Stafford, P, Higham, J, Pinnock, A, Murdoch, C, Douglas, CWI, Stafford, GP, Lambert, DW. 2013. Gingipain-dependent degradation of mammalian target of rapamycin pathway proteins by the periodontal pathogen Porphyromonas gingivalis during invasion. Molecular Oral Microbiology. 28:366–378. doi: 10.1111/omi.12030.

16. Takahara, T, Amemiya, Y, Sugiyama, R, Maki, M, Shibata, H. 2020. Amino acid-dependent control of mTORC1 signaling: a variety of regulatory modes. J Biomed Sci. 27:87. doi: 10.1186/s12929-020-00679-2.

17. Peake, J, Suzuki, K. 2004. Neutrophil activation, antioxidant supplements and exercise-induced oxidative stress. Exerc Immunol. Rev. 10:129.

18. Pakos-Zebrucka, K, Koryga, I, Mnich, K, Ljujic, M, Samali, A, Gorman, AM. 2016. The integrated stress response. EMBO Rep. 17:1374–1395. doi: 10.15252/embr.201642195.

19. Donnelly, N, Gorman, AM, Gupta, S, Samali, A. 2013. The eIF2α kinases: their structures and functions. Cell Mol Life Sci. 70:3493–3511. doi: 10.1007/s00018-012-1252-6.

20. Knowles, A, Campbell, S, Cross, N, Stafford, P. 2021. Bacterial manipulation of the Integrated Stress Response: a new perspective on infection. Front Microbiol. 12:645161.

21. Siekierka, J, Mauser, L, Ochoa, S. 1982. Mechanism of polypeptide chain initiation in eukaryotes and its control by phosphorylation of the alpha subunit of initiation factor 2. Proc Natl Acad Sci U S A. 79:2537. doi: 10.1073/pnas.79.8.2537.

22. Hinnebusch, AG, Lorsch, JR. 2012. The mechanism of eukaryotic translation initiation: new insights and challenges. Cold Spring Harbor Perspectives in Biology. 4:. doi: 10.1101/cshperspect.a011544.

23. Price, N, Proud, C. 1994. The guanine nucleotide-exchange factor, eIF-2B. Biochimie. 76:748–760. doi: 10.1016/0300-9084(94)90079-5.

24. Jennings, MD, Zhou, Y, Mohammad-Qureshi, S, Bennett, D, Pavitt, GD. 2013. eIF2B promotes eIF5 dissociation from eIF2*GDP to facilitate guanine nucleotide exchange for translation initiation. Genes Dev. 27:2696. doi: 10.1101/gad.231514.113.

25. Dever, TE, Yang, W, Aström, S, Byström, AS, Hinnebusch, AG. 1995. Modulation of tRNA(iMet), eIF-2, and eIF-2B expression shows that GCN4 translation is inversely coupled to the level of eIF-2.GTP.Met-tRNA(iMet) ternary complexes. Mol Cell Biol. 15:6351. doi: 10.1128/MCB.15.11.6351.

26. Rowlands, AG, Panniers, R, Henshaw, EC. 1988. The catalytic mechanism of guanine nucleotide exchange factor action and competitive inhibition by phosphorylated eukaryotic initiation factor 2. The Journal of Biological Chemistry. 263:5526.

27. Kenner, LR, Anand, AA, Nguyen, HC, Myasnikov, AG, Klose, CJ, Mcgeever, LA, Tsai, JC, Miller-Vedam, L, Walter, P, Frost, A. 2019. eIF2B-catalyzed nucleotide exchange and phosphoregulation by the integrated stress response. Science. 364:491. doi: 10.1126/science.aaw2922.

28. Pause, A, Belsham, GJ, Anne-Claude Gingras, Olivier Donzé, Tai-An Lin, Lawrence, JC, Sonenberg, N. 1994. Insulin-dependent stimulation of protein synthesis by phosphorylation of a regulator of 5’-cap function. Nature. 371:762. doi: 10.1038/371762a0.

29. Heberle, AM, Prentzell, MT, van Eunen, K, Bakker, BM, Grellscheid, SN, Thedieck, K. 2015. Molecular mechanisms of mTOR regulation by stress. Molecular & Cellular Oncology. 2:. doi: 10.4161/23723548.2014.970489.

30. Qin, X, Jiang, B, Zhang, Y. 2016. 4E-BP1, a multifactor regulated multifunctional protein. Cell Cycle. 15:00. doi: 10.1080/15384101.2016.1151581.

31. Nover, L, Scharf, KD, Neumann, D. 1989. Cytoplasmic heat shock granules are formed from precursor particles and are associated with a specific set of mRNAs. Mol Cell Biol. 9:1298. doi: 10.1128/MCB.9.3.1298.

32. Anderson, P, Kedersha, N. 2002. Stressful initiations. J Cell Sci. 115:3227.

33. Kedersha, N, Cho, MR, Li, W, Yacono, PW, Chen, S, Gilks, N, Golan, DE, Anderson, P. 2000. Dynamic Shuttling of Tia-1 Accompanies the Recruitment of mRNA to Mammalian Stress Granules. J Cell Biol. 151:1257–1268. doi: 10.1083/jcb.151.6.1257.

34. Loschi, M, Leishman, CC, Berardone, N, Boccaccio, GL. 2009. Dynein and kinesin regulate stress-granule and P-body dynamics. J Cell Sci. 122:3973. doi: 10.1242/jcs.051383.

35. Bussiere, LD, Miller, CL. 2021. Reovirus and the Host Integrated Stress Response: On the Frontlines of the Battle to Survive. Viruses; Viruses. 13:200. doi: 10.3390/v13020200.

36. Rabouw, HH, Visser, LJ, Passchier, TC, Langereis, MA, Liu, F, Giansanti, P, van Vliet, AL,W., Dekker, JG, van der Grein, S,G., Saucedo, JG, Anand, AA, Trellet, ME, Bonvin, Alexandre M. J. J., Walter, P, Heck, AJR, de Groot, R,J., van Kuppeveld, FJ,M. 2020. Inhibition of the integrated stress response by viral proteins that block p-eIF2–eIF2B association. Nature Microbiology. 5:1361–1373. doi: 10.1038/s41564-020-0759-0.

37. Abdel-Nour, M, Carneiro, LAM, Downey, J, Tsalikis, J, Outlioua, A, Prescott, D, Da Costa, LS, Hovingh, ES, Farahvash, A, Gaudet, RG, Molinaro, R, van Dalen, R, Lau, CCY, Azimi, FC, Escalante, NK, Trotman-Grant, A, Lee, JE, Gray-Owen, S, Divangahi, M, Chen, J, Philpott, DJ, Arnoult, D, Girardin, SE. 2019. The heme-regulated inhibitor is a cytosolic sensor of protein misfolding that controls innate immune signaling. Science. 365:. doi: 10.1126/science.aaw4144.

38. Tattoli, I, Sorbara, MT, Vuckovic, D, Ling, A, Soares, F, Carneiro, LAM, Yang, C, Emili, A, Philpott, DJ, Girardin, SE. 2012. Amino acid starvation induced by invasive bacterial pathogens triggers an innate host defense program. Cell Host & Microbe. 11:563. doi: 10.1016/j.chom.2012.04.012.

39. van ‘t Wout, Emily F.,A., van Schadewijk, A, van Boxtel, R, Dalton, LE, Clarke, HJ, Tommassen, J, Marciniak, SJ, Hiemstra, PS, Parsek, MR. 2015. Virulence Factors of Pseudomonas aeruginosa Induce Both the Unfolded Protein and Integrated Stress Responses in Airway Epithelial Cells. PLoS Pathog. 11:e1004946. doi: 10.1371/journal.ppat.1004946.

40. Smyth, R, Berton, S, Rajabalee, N, Chan, T, Sun, J. 2020. Protein Kinase R Restricts the Intracellular Survival of Mycobacterium tuberculosis by Promoting Selective Autophagy. Front Microbiol. 11:613963. doi: 10.3389/fmicb.2020.613963.

41. Shrestha, N, Boucher, J, Bahnan, W, Clark, ES, Rosqvist, R, Fields, KA, Khan, WN, Schesser, K. 2013. The host-encoded Heme Regulated Inhibitor (HRI) facilitates virulence-associated activities of bacterial pathogens. PloS One. 8:e68754. doi: 10.1371/journal.pone.0068754.

42. Tsutsuki, H, Yahiro, K, Ogura, K, Ichimura, K, Iyoda, S, Ohnishi, M, Nagasawa, S, Seto, K, Moss, J, Noda, M. 2016. Subtilase cytotoxin produced by locus of enterocyte effacement-negative Shiga-toxigenic Escherichia coli induces stress granule formation. Cell. Microbiol. 18:1024–1040. doi: 10.1111/cmi.12565.

43. Anand, A, Sharma, A, Ravins, M, Biswas, D, Ambalavanan, P, Lim, KXZ, Tan, RYM, Johri, AK, Tirosh, B, Hanski, E. 2021. Unfolded protein response inhibitors cure group A streptococcal necrotizing fasciitis by modulating host asparagine. Science Translational Medicine. 13:. doi: 10.1126/scitranslmed.abd7465.

44. Velásquez, F, Marín-Rojas, J, Soto-Rifo, R, Torres, A, Del Canto, F, Valiente-Echeverría, F. 2020. Escherichia coli HS and Enterotoxigenic Escherichia coli Hinder Stress Granule Assembly. Microorganisms. 9:17. doi: 10.3390/microorganisms9010017.

45. Vonaesch, P, Campbell-Valois, F, Dufour, A, Sansonetti, PJ, Schnupf, P. 2016. Shigella flexneri modulates stress granule composition and inhibits stress granule aggregation. Cell Microbiol. 18:982–997. doi: 10.1111/cmi.12561.

46. Nakayama, M, Inoue, T, Naito, M, Nakayama, K, Ohara, N. 2015. Attenuation of the Phosphatidylinositol 3-Kinase/Akt Signaling Pathway by Porphyromonas gingivalis Gingipains RgpA, RgpB, and Kgp. J Biol Chem. 290:5190–5202. doi: 10.1074/jbc.M114.591610.

47. Hirasawa, M, Kurita-Ochiai, T. 2018. Porphyromonas gingivalis Induces Apoptosis and Autophagy via ER Stress in Human Umbilical Vein Endothelial Cells. Mediators Inflamm. 2018:1967506–8. doi: 10.1155/2018/1967506.

48. Harding, HP, Zhang, Y, Zeng, H, Novoa, I, Lu, PD, Calfon, M, Sadri, N, Yun, C, Popko, B, Paules, R, Stojdl, DF, Bell, JC, Hettmann, T, Leiden, JM, Ron, D. 2003. An Integrated Stress Response Regulates Amino Acid Metabolism and Resistance to Oxidative Stress. Mol Cell. 11:619–633. doi: 10.1016/S1097-2765(03)00105-9.

49. Aulas, A, Fay, MM, Lyons, SM, Achorn, CA, Kedersha, N, Anderson, P, Ivanov, P. 2017. Stress-specific differences in assembly and composition of stress granules and related foci. J Cell Sci. 130:927–937. doi: 10.1242/jcs.199240.

50. Sidrauski, C, McGeachy, AM, Ingolia, NT, Walter, P. 2015. The small molecule ISRIB reverses the effects of eIF2α phosphorylation on translation and stress granule assembly. eLife. 4:. doi: 10.7554/eLife.05033.

51. Nandagopal, N, Roux, PP. 2015. Regulation of global and specific mRNA translation by the mTOR signaling pathway. Translation. 3:e983402. doi: 10.4161/21690731.2014.983402.

52. Wheeler, JR, Matheny, T, Jain, S, Abrisch, R, Parker, R. 2016. Distinct stages in stress granule assembly and disassembly. eLife. 5:. doi: 10.7554/eLife.18413.

53. Costa-Mattioli, M, Walter, P. 2020. The integrated stress response: From mechanism to disease. Science. 368:. doi: 10.1126/science.aat5314.

54. Pavitt, GD, Ron, D. 2012. New insights into translational regulation in the endoplasmic reticulum unfolded protein response. Cold Spring Harb Perspect Biol. 4:a012278. doi: 10.1101/cshperspect.a012278.

55. Mayadas, TN, Cullere, X, Lowell, CA. 2014. The multifaceted functions of neutrophils. Annu Rev Pathol. 9:181–218. doi: 10.1146/annurev-pathol-020712-164023.

56. McEwen, E, Kedersha, N, Song, B, Scheuner, D, Gilks, N, Han, A, Chen, J, Anderson, P, Kaufman, RJ. 2005. Heme-regulated Inhibitor Kinase-mediated Phosphorylation of Eukaryotic Translation Initiation Factor 2 Inhibits Translation, Induces Stress Granule Formation, and Mediates Survival upon Arsenite Exposure. J Biol Chem. 280:16925–16933. doi: 10.1074/jbc.M412882200.

57. Cekici, A, Kantarci, A, Hasturk, H, Van Dyke, T,E. 2014. Inflammatory and immune pathways in the pathogenesis of periodontal disease: Inflammatory and immune pathways in periodontal disease. Periodontol 2000. 64:57–80. doi: 10.1111/prd.12002.

58. Henry, LG, McKenzie, RME, Robles, A, Fletcher, HM. 2012. Oxidative stress resistance in Porphyromonas gingivalis. Future Microbiol. 7:497–512. doi: 10.2217/fmb.12.17.

59. Choi, CH, Spooner, R, DeGuzman, J, Koutouzis, T, Ojcius, DM, Yilmaz, Ö. 2013. P. gingivalis-Nucleoside-diphosphate-kinase Inhibits ATP-Induced Reactive-Oxygen-Species via P2X7 Receptor/NADPH-Oxidase Signaling and Contributes to Persistence. Cell Microbiol. 15:961–976. doi: 10.1111/cmi.12089.

60. Smalley, JW, Birss, AJ, Silver, J. 2000. The periodontal pathogen Porphyromonas gingivalis harnesses the chemistry of the µ-oxo bishaem of iron protoporphyrin IX to protect against hydrogen peroxide. FEMS Microbiol Lett. 183:159–164. doi: 10.1016/S0378-1097(99)00660-6.

61. Johnson, NA, Mckenzie, R, Mclean, L, Sowers, LC, Fletcher, HM. 2004. 8-Oxo-7,8-Dihydroguanine Is Removed by a Nucleotide Excision Repair-Like Mechanism in Porphyromonas gingivalis W83. J Bacteriol. 186:7697–7703. doi: 10.1128/JB.186.22.7697-7703.2004.

62. Yancy, SL, Shelden, EA, Gilmont, RR, Welsh, MJ. 2005. Sodium Arsenite Exposure Alters Cell Migration, Focal Adhesion Localization and Decreases Tyrosine Phosphorylation of Focal Adhesion Kinase in H9C2 Myoblasts. Toxicol Sci. 84:278–286. doi: 10.1093/toxsci/kfi032.

63. Fang, MY, Markmiller, S, Vu, AQ, Javaherian, A, Dowdle, WE, Jolivet, P, Bushway, PJ, Castello, NA, Baral, A, Chan, MY, Linsley, JW, Linsley, D, Mercola, M, Finkbeiner, S, Lecuyer, E, Lewcock, JW, Yeo, GW. 2019. Small-Molecule Modulation of TDP-43 Recruitment to Stress Granules Prevents Persistent TDP-43 Accumulation in ALS/FTD. Neuron. 103:802–819.e11. doi: 10.1016/j.neuron.2019.05.048.

64. Gingras, A, Raught, B, Sonenberg, N. 1999. eIF4 INITIATION FACTORS: Effectors of mRNA Recruitment to Ribosomes and Regulators of Translation. Annu Rev Biochem. 68:913–963. doi: 10.1146/annurev.biochem.68.1.913.

65. Sévigny, M, Bourdeau Julien, I, Venkatasubramani, JP, Hui, JB, Dutchak, PA, Sephton, CF. 2020. FUS contributes to mTOR-dependent inhibition of translation. J Biol Chem. 295:18459–18473. doi: 10.1074/jbc.RA120.013801.

66. Larsson, O, Morita, M, Topisirovic, I, Alain, T, Blouin, M, Pollak, M, Sonenberg, N. 2012. Distinct perturbation of the translatome by the antidiabetic drug metformin. Proc Natl Acad Sci U S A. 109:8977–8982. doi: 10.1073/pnas.1201689109.

67. Holz, MK, Ballif, BA, Gygi, SP, Blenis, J. 2005. mTOR and S6K1 Mediate Assembly of the Translation Preinitiation Complex through Dynamic Protein Interchange and Ordered Phosphorylation Events. Cell. 123:569–580. doi: 10.1016/j.cell.2005.10.024.

68. Wu, X, Wang, X, Wu, S, Lu, J, Zheng, M, Wang, Y, Zhou, H, Zhang, H, Han, J. 2011. Phosphorylation of Raptor by p38β Participates in Arsenite-induced Mammalian Target of Rapamycin Complex 1 (mTORC1) Activation. J Biol Chem. 286:31501–31511. doi: 10.1074/jbc.M111.233122.

69. Vasquez, RJ, Howell, B, Yvon, AM, Wadsworth, P, Cassimeris, L. 1997. Nanomolar concentrations of nocodazole alter microtubule dynamic instability in vivo and in vitro. Mol Biol Cell. 8:973–985. doi: 10.1091/mbc.8.6.973.

70. Nadezhdina, ES, Lomakin, AJ, Shpilman, AA, Chudinova, EM, Ivanov, PA. 2010. Microtubules govern stress granule mobility and dynamics. BBA - Molecular Cell Research. 1803:361–371. doi: 10.1016/j.bbamcr.2009.12.004.

71. Kinane, JA, Benakanakere, MR, Zhao, J, Hosur, KB, Kinane, DF. 2012. Porphyromonas gingivalis influences actin degradation within epithelial cells during invasion and apoptosis. Cell Microbiol. 14:1085–1096. doi: 10.1111/j.1462-5822.2012.01780.x.

72. Magiera, MM, Janke, C. 2014. Post-translational modifications of tubulin. Current Biology. 24:R351–R354. doi: 10.1016/j.cub.2014.03.032.

73. Reed, NA, Cai, D, Blasius, TL, Jih, GT, Meyhofer, E, Gaertig, J, Verhey, KJ. 2006. Microtubule acetylation promotes kinesin-1 binding and transport. Current Biology. 16:2166.

74. Cai, D, McEwen, DP, Martens, JR, Meyhofer, E, Verhey, KJ, Schliwa, M. 2009. Single Molecule Imaging Reveals Differences in Microtubule Track Selection Between Kinesin Motors. PLoS Biology. 7:e1000216. doi: 10.1371/journal.pbio.1000216.

75. Hammond, JW, Huang, C, Kaech, S, Jacobson, C, Banker, G, Verhey, KJ, Holzbaur, E. 2010. Posttranslational Modifications of Tubulin and the Polarized Transport of Kinesin-1 in Neurons. Mol Biol Cell. 21:572–583. doi: 10.1091/mbc.E09-01-0044.

76. Zhang, Y, Li, N, Caron, C, Matthias, G, Hess, D, Khochbin, S, Matthias, P. 2003. HDAC-6 interacts with and deacetylates tubulin and microtubules in vivo. Embo J. 22:1168–1179. doi: 10.1093/emboj/cdg115.

77. Kwon, S, Zhang, Y, Matthias, P. 2007. The deacetylase HDAC6 is a novel critical component of stress granules involved in the stress response. Genes Dev. 21:3381–3394. doi: 10.1101/gad.461107.

78. Liu, Y, Wang, M, Cheng, A, Yang, Q, Wu, Y, Jia, R, Liu, M, Zhu, D, Chen, S, Zhang, S, Zhao, X, Huang, J, Mao, S, Ou, X, Gao, Q, Wang, Y, Xu, Z, Chen, Z, Zhu, L, Luo, Q, Liu, Y, Yu, Y, Zhang, L, Tian, B, Pan, L, Rehman, MU, Chen, X. 2020. The role of host eIF2α in viral infection. Virol J. 17:112. doi: 10.1186/s12985-020-01362-6.

79. Zyryanova, AF, Kashiwagi, K, Rato, C, Harding, HP, Crespillo-Casado, A, Perera, LA, Sakamoto, A, Nishimoto, M, Yonemochi, M, Shirouzu, M, Ito, T, Ron, D. 2021. ISRIB Blunts the Integrated Stress Response by Allosterically Antagonising the Inhibitory Effect of Phosphorylated eIF2 on eIF2B. Mol Cell. 81:88–103.e6. doi: 10.1016/j.molcel.2020.10.031.

80. Schoof, M, Boone, M, Wang, L, Lawrence, R, Frost, A, Walter, P. 2021. eIF2B Conformation and Assembly State Regulate the Integrated Stress Response. The FASEB Journal. 35:e65703. doi: 10.1096/fasebj.2021.35.S1.04030.

81. Potempa, J, Travis, J. 1996. Porphyromonas gingivalis proteinases in periodontitis, a review. Acta Biochim. Pol. 43:455–465. doi: 10.18388/abp.1996_4477.

82. Potempa, J, Pike, R, Travis, J. 1995. The multiple forms of trypsin-like activity present in various strains of Porphyromonas gingivalis are due to the presence of either Arg-gingipain or Lys-gingipain. Infect Immun. 63:1176–1182. doi: 10.1128/IAI.63.4.1176-1182.1995.

83. Mantri, CK, Chen, C, Dong, X, Goodwin, JS, Pratap, S, Paromov, V, Xie, H. 2015. Fimbriae-mediated outer membrane vesicle production and invasion of Porphyromonas gingivalis. MicrobiologyOpen. 4:53–65. doi: 10.1002/mbo3.221.

84. Xia, Q, Wang, T, Taub, F, Park, Y, Capestany, CA, Lamont, RJ, Hackett, M. 2007. Quantitative proteomics of intracellular Porphyromonas gingivalis. Proteomics. 7:4323–4337. doi: 10.1002/pmic.200700543.

85. Wolozin, B, Ivanov, P. 2019. Stress granules and neurodegeneration. Nat Rev Neurosci. 20:649–666. doi: 10.1038/s41583-019-0222-5.

86. Vaklavas, C, Blume, SW, Grizzle, WE. 2017. Translational Dysregulation in Cancer: Molecular Insights and Potential Clinical Applications in Biomarker Development. Front Oncol. 7:158. doi: 10.3389/fonc.2017.00158.

87. Aduse-Opoku, J, Davies, NN, Gallagher, A, Hashim, A, Evans, HEA, Rangarajan, M, Slaney, JM, Curtis, MA. 2000. Generation of Lys-gingipain protease activity in Porphyromonas gingivalis W50 is independent of Arg-gingipain protease activity. Microbiology. 146:1933–1940. doi: 10.1099/00221287-146-8-1933.

88. Dong, X, Ho, M, Liu, B, Hildreth, J, Dash, C, Goodwin, JS, Balasubramaniam, M, Chen, C, Xie, H. 2018. Role of Porphyromonas gingivalis outer membrane vesicles in oral mucosal transmission of HIV. Sci Rep. 8:8812–10. doi: 10.1038/s41598-018-27284-6.

89. Hodgson, RE, Varanda, BA, Ashe, MP, Allen, KE, Campbell, SG. 2019. Cellular eIF2B subunit localization: implications for the integrated stress response and its control by small molecule drugs. Mol Biol Cell. 30:942–958. doi: 10.1091/mbc.E18-08-0538.

